# *Klebsiella pneumoniae* reduces SUMOylation to limit host defence responses

**DOI:** 10.1101/2020.06.29.179275

**Authors:** Joana Sá-Pessoa, Kornelia Przybyszewska, Filipe Nuno Vasconcelos, Amy Dumigan, Christian G. Frank, Laura Hobley, Jose A. Bengoechea

## Abstract

*Klebsiella pneumoniae* is an important cause of multidrug resistant infections worldwide. Understanding the virulence mechanisms of *K. pneumoniae* is a priority and timely to design new therapeutics. Here we demonstrate that *K. pneumoniae* limits the SUMOylation of host proteins in epithelial cells and macrophages (mouse and human) to subvert cell innate immunity. Mechanistically, in lung epithelial cells *Klebsiella* increases the levels of the deSUMOylase SENP2 in the cytosol by affecting its K48-ubiquitylation and its subsequent degradation by the ubiquitin proteasome. This is dependent on *Klebsiella* preventing the NEDDylation of the Cullin-1 subunit of the ubiquitin ligase complex E3-SCFβ-TrCP by exploiting the CSN5 deNEDDylase. *Klebsiella* induces the expression of CSN5 in an EGFR-PI3K-AKT-ERK-GSK3β signalling pathway dependent manner. In macrophages, TLR4-TRAM-TRIF induced type-I IFN via IFNAR1-controlled signalling mediates *Klebsiella*-triggered decrease in the levels of SUMOylation via *let-7* miRNAs. Our results revealed the crucial role played by *Klebsiella* polysaccharides, the capsule and the LPS O-polysaccharide, to decrease the levels of SUMO-conjugated proteins in epithelial cells and macrophages. *Klebsiella*-induced decrease in SUMOylation promotes infection by limiting the activation of inflammatory responses and increasing intracellular survival in macrophages.

**IMPORTANCE:** *Klebsiella pneumoniae* has been singled out as an urgent threat to human health due to the increasing isolation of strains resistant to “last line” antimicrobials, narrowing the treatment options against *Klebsiella* infections. Unfortunately, at present, we cannot identify candidate compounds in late-stage development for treatment of multidrug *Klebsiella* infections; this pathogen is exemplary of the mismatch between unmet medical needs and the current antimicrobial research and development pipeline. Furthermore, there is still limited evidence on *K. pneumoniae* pathogenesis at the molecular and cellular level in the context of the interactions between bacterial pathogens and their hosts. In this research, we have uncovered a sophisticated strategy employed by *Klebsiella* to subvert the activation of immune defences by controlling the modification of proteins. Our research may open opportunities to develop new therapeutics based on counteracting this *Klebsiella-*controlled immune evasion strategy.

## INTRODUCTION

Posttranslational modifications (PTMs) are unique mechanisms that allow cells to rapidly and specifically modify activity or interactions of proteins in a reversible process. SUMOylation is a crucial PTM involved in cell cycle, metabolism, stress response, transcription regulation, and many other biological processes. Proteomic studies have uncovered that ∼1,000 proteins are SUMO targets [1]. Modification of proteins by SUMO is achieved through activation of SUMO by an E1 activating enzyme (the heterodimer SAE1/SAE2); transfer of this activated SUMO to an E2 conjugating enzyme (Ubc9); and ligation to the substrates catalysed by different E3 ligases: members of the PIAS (Protein Inhibitors of Activated STAT) family, RanBP2, members of the Polycomb group proteins, and of the TRIM (tripartite motif) family [2]. In parallel, deconjugation of SUMO is carried by either DeSI enzymes (DeSumoylating Isopeptidases, DeSI-1 and DeSI-2) [3] or a family of six sentric-specific proteases (SENPs) that cleave the inactive form of SUMO starting the SUMOylation cycle and have isopeptidase activity recycling SUMO from substrate proteins [3].

It is not surprising that pathogens target SUMOylation given its importance in controlling many pathways in eukaryotic cells. There is a wealth of evidence demonstrating that SUMOylation is involved in many aspects of the cell-virus interplay, either promoting or interfering with infection [4]. In contrast, the interplay between bacterial pathogens and host SUMOylation is less well understood. To date, it has been shown that the intracellular pathogens *Listeria monocytogenes, Shigella* and *Salmonella typhimurium* trigger an overall decrease in SUMOylation of proteins [5–7]. The three pathogens induce the degradation of the E2 enzyme Ubc9 to decrease the amount of SUMO conjugates [5-7], indicating that depletion of Ubc9 could be a preferred bacterial strategy to target SUMOylation. Recently, *S. flexneri* has also been shown to trigger the degradation of the SUMO E1 enzyme SAE2 to reduce SUMOylation [8].

We decided to study the interplay between SUMOylation and the human pathogen *Klebsiella pneumoniae*. This Gram-negative bacterium causes a wide range of infections, from urinary tract infections to pneumonia. The latter is particularly devastating among immunocompromised patients. *K. pneumoniae* is a member of the so-called ESKAPE group of microorganisms to emphasise that they effectively “escape” the effects of antibacterial drugs. Therefore, the development of new therapeutic strategies requires a better understanding of *K. pneumoniae* biology in the context of the complex interactions between bacterial pathogens and their hosts. This pathogens has developed sophisticated strategies to attenuate the activation of host defence [9], and therefore, we hypothesized that *Klebsiella* may target SUMOylation to promote infection. A wealth of evidence underscores the importance of the Toll-like receptor (TLR)-governed inflammatory response to clear *K. pneumoniae* infections [10,11]. In turn, we and others have provided compelling evidence demonstrating that one *Klebsiella* virulence strategy is the evasion of TLR2/4-controlled antimicrobial defences [12,13]. *Klebsiella* manipulates TLR4, EGF receptor and NOD1 signalling to ablate the activation of NF-κB and mitogen-activated protein kinases (MAPKs) [14,15].

Here, we demonstrate that *K. pneumoniae* impairs the SUMOylation of host proteins in epithelial cells and macrophages to subvert cell innate immunity. Mechanistically, *Klebsiella* utilises different strategies depending on the type of cell. In epithelial cells, *Klebsiella* exploits the SENP2 deSUMOylase by preventing its degradation by the ubiquitin proteasome whereas in macrophages the decrease in SUMOylated proteins is dependent on a type-I IFN-induced micro RNA (miRNA) of the *let-7* family.

## RESULTS

### *K. pneumoniae* decreases SUMO-conjugated proteins in epithelial cells

To investigate whether *K. pneumoniae* affects host cell SUMOylation, we compared the global pattern of proteins conjugated to SUMO1 or SUMO2/3 in uninfected cells with that of cells infected by the wild-type hypervirulent *K. pneumoniae* strain CIP52.145 (Kp52145 hereafter). This strain encodes the genetic determinants associated with severe human infections [16]. A549 epithelial cells showed a reduction in overall SUMO1 conjugated proteins of high molecular weight (above 80 kDa) after 3h of infection with no changes in the overall SUMO2/3 conjugated proteins (Figure 1A). We then validated the decrease in SUMOylation on the heavily SUMOylated RanGAP1 substrate. In *Klebsiella*-infected cells, we observed a clear reduction in SUMOylated RanGAP1 after 3 hours post infection (Figure 1B). The decrease in SUMO1 conjugated proteins was also observed in NuLi-1 cells, a human primary-like airway epithelial cell line (Supplementary Figure S1), demonstrating that *Klebsiella*-induced decrease in overall SUMOylation is not cell type dependent. Kp52145-induced decrease in protein SUMOylation was dependent on live bacteria because infection with either UV-killed or heat-killed bacteria caused an increase in SUMO1 conjugated proteins (Figure 1C). To examine the relevance of these findings during infection, we assessed the pattern of SUMOylated proteins in lungs of infected mice. Although there was mouse to mouse variation, immunoblotting of lung homogenates revealed a decrease in the global SUMO1-conjugated proteome compared to homogenates from mock-infected mice (Figure 1D). Together, our *in vitro* and *in vivo* findings reveal that *K. pneumoniae* infection leads to a decrease in the levels of SUMO-conjugated proteins.

**Figure 1.**
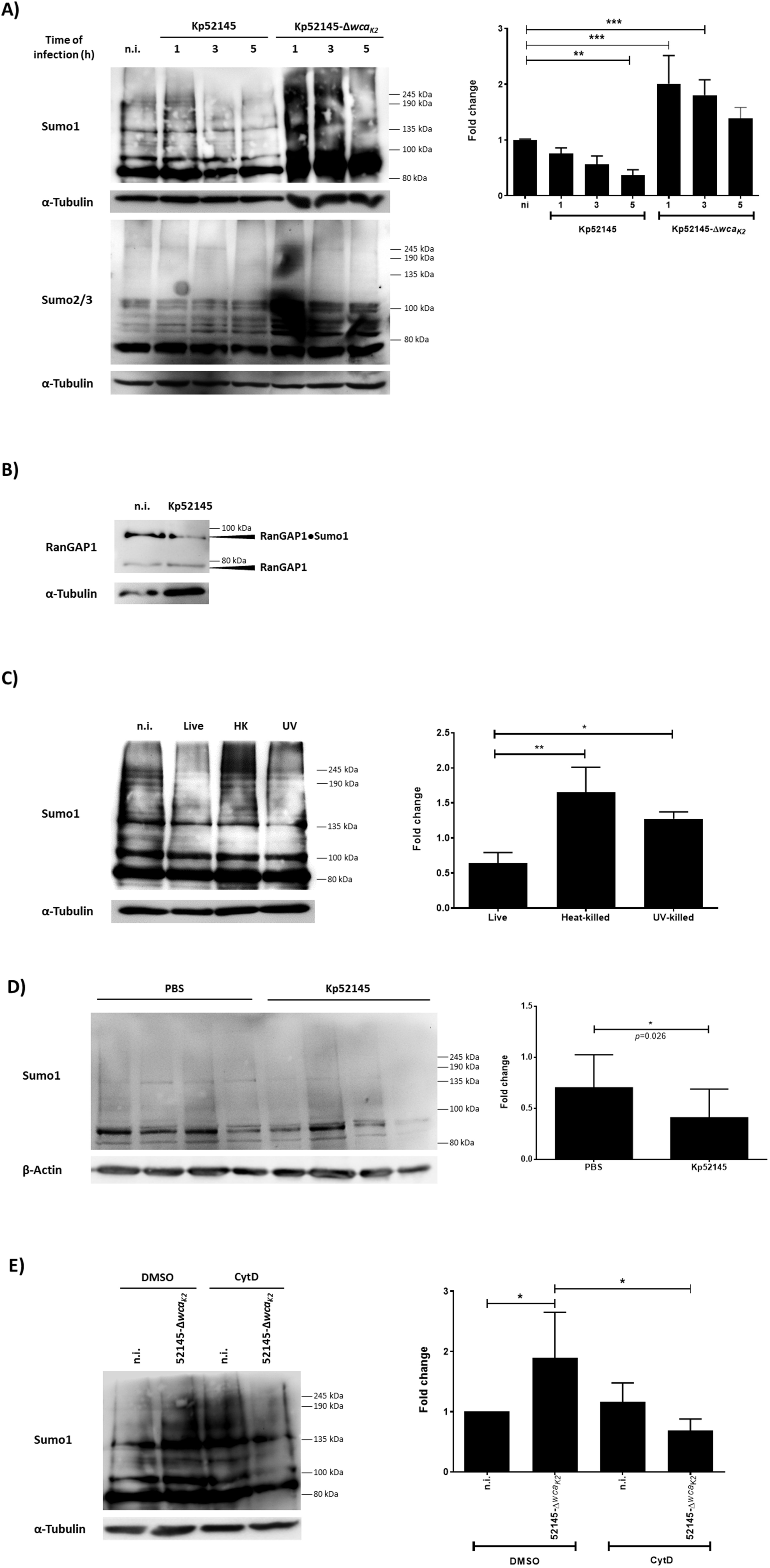
*Klebsiella pneumoniae* decreases SUMO-conjugated proteins in epithelial cells. A. Immunoblot analysis of SUMO1, SUMO2/3 and tubulin levels in lysates of A549 cells infected with Kp52145 and the capsule mutant (strain 52145-Δ*wca*_*K2*_) for the indicated times. SUMO1 smears were quantified from three independent experiments using Image Studio Lite (Li-cor) and the modification normalised to α-Tubulin. The graph represents fold change compared to non-infected control cells. **P ≤ 0.01; *P ≤ 0.05; versus n.i. determined using one way-ANOVA with Bonferroni’s multiple comparisons test. B. Immunoblot analysis of RanGAP1 and tubulin levels in A549 infected with Kp52145 for the indicated times. C. Immunoblot analysis of SUMO1 and tubulin levels in lysates of A549 cells infected with Kp52145 live, heat-killed for 30 min at 60 °C or UV-killed for 30 min at 10000 J for the indicated times. SUMO1 smears were quantified from three independent experiments using Image Studio Lite (Li-cor) and the modification normalised to α-Tubulin. The graph represents fold change at 5 h of infection compared to non-infected control cells. **P ≤ 0.01; *P ≤ 0.05; versus n.i. determined using one way-ANOVA with Bonferroni’s multiple comparisons test. D.Immunoblot analysis of SUMO1 and tubulin levels in 50 µg of total protein obtained from whole lung homogenate of C57BL/6 mice infected with Kp52145 or with PBS as a mock control for 24 h. Each lane represents a lung from a different mouse. SUMO1 smears were quantified from three independent blots using Image Studio Lite (Li-cor) and the modification normalised to β-actin. The graph represents fold change of the Kp52145 lung homogenates compared to PBS controls. *P = 0.0265 versus PBS determined using a two-tailed unpaired *t*-test. E.Immunoblot analysis of SUMO1 levels in lysates of cytochalasin D (CytD, 5 μM, 1 h before infection) or DMSO (vehicle solution)-treated A549 cells infected with strain Kp52145-Δ*wca*_*K2*_ for 5 h. Membranes were reprobed for tubulin as a loading control. SUMO1 smears were quantified from three independent experiments using Image Studio Lite (Li-cor) and the modification normalised to α-Tubulin. The graph represents fold change compared to DMSO non-infected control cells. **P ≤ 0.01; *P ≤ 0.05; versus DMSO n.i. determined using one way-ANOVA with Bonferroni’s multiple comparisons test. n.i. – non-infected control In all panels data are representative of at least three independent experiments.

In contrast to the wild-type strain, infection with an isogenic avirulent capsule mutant (strain 52145-Δ*wca*_*K2*_) induced an increase in SUMO1 and SUMO2/3 conjugated proteins (Figure 1A). Since the capsule polysaccharide (CPS) abrogates the internalisation of *K. pneumoniae* by A549 [17], we hypothesised that the capsule mutant-induced increase in SUMOylation is triggered by bacterial internalisation. Indeed, when infections were performed in the presence of cytochalasin D, an inhibitor of *Klebsiella* internalisation by epithelial cells [17], the CPS mutant did not increase the levels of SUMO1-conjugated proteins (Figure 1E). Collectively, these results indicate that the CPS abrogates the increase in SUMOylation upon *Klebsiella* infection by limiting the pathogen internalization.

### The deSUMOylase SENP2 mediates *K. pneumoniae*-induced decrease in SUMO1-conjugated proteins in epithelial cells

We sought to gain insights into the mechanisms by which wild-type *Klebsiella* reduces the SUMOylation status in epithelial cells. Previous studies indicate that depletion of the E1 enzyme SAE2, and the E2 enzyme Ubc9 underlines bacterial pathogen-triggered decrease in SUMOylation [5–8]. However, the levels of Ubc9 were not significantly affected in Kp52145-infected cells (Figure 2A), and we did not detect any change in the levels of SAE2 or SAE1, another E1 enzyme (Figure 2B). Although these results do not rigorously rule out that *Klebsiella* may affect the enzymatic activity of these enzymes, it prompted us to consider alternative possibilities to explain the *Klebsiella*-induced decrease in SUMOylation.

**Figure 2.**
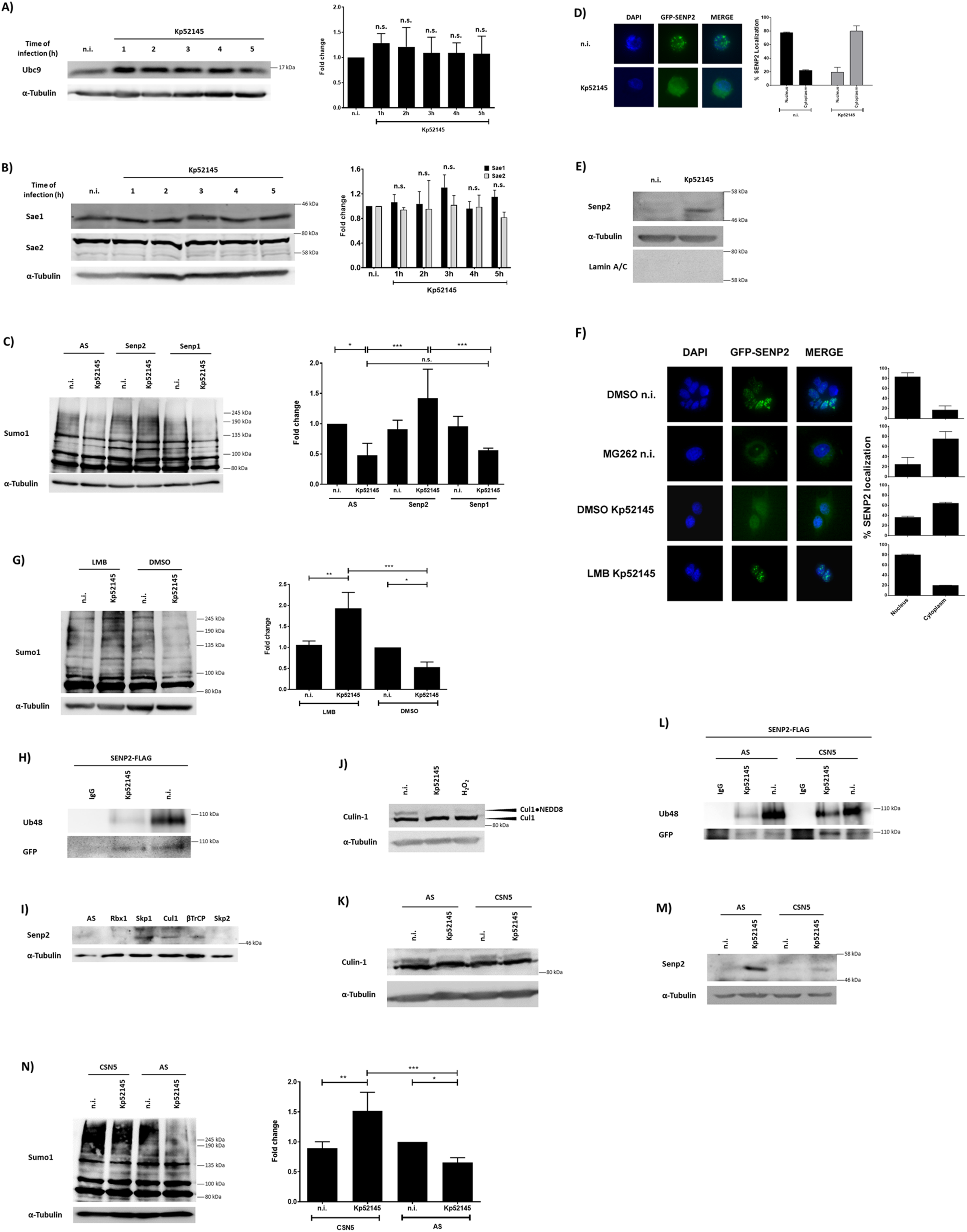
The deSUMOylase SENP2 mediates *K. pneumoniae*-induced decrease in SUMO-conjugated proteins. A. Immunoblot analysis of Ubc9 and tubulin levels in lysates of A549 cells infected with Kp52145 for the indicated times. Ubc9 bands were quantified from three independent experiments using Image Studio Lite (Li-cor) and the modification normalised to α-Tubulin. The graph represents fold change compared to non-infected control cells. n.s. – non significant; versus n.i. determined using one way-ANOVA with Tukey’s multiple comparisons test. B. Immunoblot analysis of Sae1, Sae2 and tubulin levels in lysates of A549 cells infected with Kp52145 for the indicated times. Sae1 and Sae2 bands were quantified from three independent experiments using Image Studio Lite (Li-cor) and the modification normalised to α-Tubulin. The graph represents fold change compared to non-infected control cells. n.s. – non significant; versus n.i. determined using one way-ANOVA with Tukey’s multiple comparisons test. C. Immunoblot analysis of SUMO1 and tubulin levels in lysates of control (AS), SENP1 or SENP2 siRNA-transfected A549 cells infected with Kp52145 for 5 h. AS-AllStars control, non-silencing siRNA. SUMO1 smears were quantified from three independent experiments using Image Studio Lite (Li-cor) and the modification normalised to α-Tubulin. The graph represents fold change compared to AS-transfected non-infected control cells. ***P ≤ 0.001; **P ≤ 0.01; *P ≤ 0.05; versus AS n.i. determined using one way-ANOVA with Bonferroni’s multiple comparisons test. D.Fluorescence microscopy of pSENP2-GFP transfected A549 cells grown on glass coverslips. Cells were infected with Kp52145 for 5 h or left uninfected (n.i.). Coverslips were stained with Hoechst (DAPI) for nuclei identification. The % of SENP2 localised in either nucleus or cytoplasm is represented on the graph and is the result of independent counting of 100 cells from each of 3 independent experiments. E.Immunoblot analysis of SENP2 and tubulin levels in cytosolic extracts of A549 cells infected with Kp52145 for 5 h or left uninfected (n.i.). F.Fluorescence microscopy of pSENP2-GFP transfected A549 cells grown on glass coverslips. Cells were treated with the proteasomal inhibitor MG262 (5 μM, 2 h before infection), the nuclear export inhibitor Leptomycin B (LMB, 10 nM, 2 h before infection) or DMSO (vehicle solution) and infected with Kp52145 for 5 h or left uninfected (n.i.). Coverslips were stained with Hoechst (DAPI) for nuclei identification. The % of SENP2 localized in either nucleus or cytoplasm is represented on the graphs and is the result of independent counting of 100 cells from each of 3 independent experiments. G.Immunoblot analysis of SUMO1 and tubulin levels in lysates of the nuclear export inhibitor Leptomycin B (LMB, 5 μM, 2 h before infection) or DMSO (vehicle solution)-treated A549 infected with Kp52145 for 5 h or left uninfected (n.i.). SUMO1 smears were quantified using Image Studio Lite (Li-cor) and the modification normalised to α-Tubulin. The graph represents fold change compared to DMSO non-infected control cells. **P ≤ 0.01 versus DMSO n.i. determined using one way-ANOVA with Bonferroni’s multiple comparisons test. H.Immunoblot analysis of K48-linkage specific polyubiquitin and FLAG (DYKDDDDK-protein tag) levels in immunoprecipitates of A549 infected with Kp52145 for 5 h. Cells were immunoprecipitated using anti-FLAG antibody. Preimmune mouse IgG served as negative control. I.Immunoblot analysis of SENP2 and tubulin levels in cytosolic extracts of control (AS), Rbx1, Skp1, Cul-1, βTrCP or Skp2 siRNA-transfected A549 cells left uninfected (n.i.). J.Immunoblot analysis of Cullin-1 and tubulin levels in lysates of A549 cells infected with Kp52145 for 5 h or left uninfected (n.i.). Lysates of A549 treated with 5 mM of H_2_O_2_ for 10 min were used as a positive control for Cullin-1 deNEDDylation. K.Immunoblot analysis of Cullin-1 and tubulin levels in lysates of control (AS) or CSN5 siRNA-transfected A549 cells infected with Kp52145 for 5 h or left uninfected (n.i.). AS-AllStars control, non-silencing siRNA. L.Immunoblot analysis of K48-linkage specific polyubiquitin and FLAG (DYKDDDDK-protein tag) levels in immunoprecipitates of control (AS) or CSN5 siRNA-transfected A549. Cells were transfected with a SENP2-FLAG plasmid 24 h after the siRNA transfection and infected the following day with Kp52145 for 5 h or left uninfected. Cells were immunoprecipitated using anti-FLAG antibody. Preimmune mouse IgG served as negative control. M.Immunoblot analysis of Senp2 and tubulin levels in cytosolic extracts of control (AS) or CSN5 siRNA-transfected A549 cells infected with Kp52145 for 5 h or left uninfected (n.i.). AS-AllStars control, non-silencing siRNA. N.Immunoblot analysis of SUMO1 and tubulin levels in lysates of control (AS) or CSN5 siRNA-transfected A549 cells infected with Kp52145 for 5 h or left uninfected (n.i.). AS-AllStars control, non-silencing siRNA. SUMO1 smears were quantified from three independent experiments using Image Studio Lite (Li-cor) and the modification normalised to α-Tubulin. The graph represents fold change compared to AS-transfected non-infected control cells. **P ≤ 0.01 versus AS n.i. determined using one way-ANOVA with Bonferroni’s multiple comparisons test. In all panels data are representative of at least three independent experiments.

SENPs, a family of cysteine proteases, catalyse the deconjugation of SUMO from target substrates; therefore, we hypothesised that they might be involved in *Klebsiella*-triggered reduction of SUMO-conjugated proteins. SENP1 and SENP2 have nuclear localisation with isopeptidase and hydrolase activity for SUMO1 and SUMO2/3 [18]. SENP3 and SENP5 are localised in the nucleolus while SENP6 and SENP7 are localised in the nucleoplasm and all favour SUMO2/3 as substrates [18]. An siRNA-based screen revealed an increase in the levels of SUMO1-conjugated proteins in infected SENP2 knockdown cells compared to cells transfected with non-silencing control siRNA (AS) and the other SENPs (Figure 2C, Supplementary Figure S2A), suggesting that SENP2 mediates the decrease in conjugated SUMO1 proteins in *K. pneumoniae* infected cells. Control experiments showed that *Klebsiella* infection did not alter the transcription of any SENPs (Supplementary Figure S2B).

Owing to the global scale of SUMOylation decrease in infected cells, it was intriguing that SENP2, a nuclear associated deSUMOylase [19], accounts for the *Klebsiella*-triggered decrease in SUMOylation. We then sought to determine whether *Klebsiella* might affect the cellular distribution of SENP2 in infected cells. Fluorescence-based single-cell analysis confirmed that SENP2-GFP was mostly localised in the nuclei of non-infected A549 cells (Figure 2D). In contrast, SENP2-GFP was mostly localised in the cytosol of infected cells (Figure 2D). This finding was further supported by immunoblotting of cytosolic extracts demonstrating an accumulation of endogenous SENP2 in infected cells (Figure 2E).

In human bone osteosarcoma epithelial cells (U-2 OS), it has been shown that SENP2 shuttles continuously between the nucleus and the cytoplasm where it is degraded by the 26S ubiquitin proteasome [20]. Corroborating these findings, pre-treatment of A549 cells with the proteasome inhibitor MG262 led to the accumulation of SENP2-GFP in the cytosol (Figure 2F). The localisation of SENP2-GFP in the cytosol of infected cells was abrogated when infections were performed in the presence of leptomycin B, a nuclear export inhibitor (Figure 2F), demonstrating that the presence of SENP2 in the cytosol of infected cells is dependent on nucleo-cytoplasmic shuttling. The *Klebsiella*-induced decrease in SUMOylation was not observed in leptomycin B-treated cells (Figure 2G), indicating that *Klebsiella*-triggered SENP2 localisation in the cytosol mediates the decrease in the levels of SUMO-conjugated proteins.

### *K. pneumoniae* triggers loss of Cullin-1 NEDDylation to increase SENP2 levels

To gain molecular insights into how *Klebsiella* increases the levels of SENP2 in the cytosol of infected cells, we hypothesized that *K. pneumoniae* may compromise the ubiquitin proteasome-mediated degradation of SENP2 occurring in the cytosol [20]. The fact that SENP2 ubiquitylation is a necessary event to mark the protein for degradation by the 26S proteasome [20] prompted us to assess whether SENP2 is ubiquitylated in infected cells. In MG262-treated cells, FLAG-tagged SENP2 was pulled down with an anti-FLAG antibody and K48-conjugated SENP2 was detected with an anti-K48 antibody. Ubiquitylated SENP2 was barely detected in *Klebsiella* infected cells in contrast to non-infected cells (Figure 2H), sustaining the notion that SENP2 accumulation in the cytosol of infected cells is due to reduced proteasomal degradation because SENP2 is not K48 ubiquitin-tagged.

We next sought to identify the E3 ubiquitin ligase complex responsible for SENP2 ubiquitylation. The SCF (SKP1 (S-phase-kinase-associated protein 1), Cullin-1, F-box protein) E3 ubiquitin ligases are the largest family of E3s in mammals [21]. Reduction in the levels of the adaptor protein Skp1 and the scaffold protein Cullin-1 (Cul-1) by siRNA led to an accumulation of SENP2 in the cytosol of non-infected cells (Figure 2I). The F-box protein determines the substrate specificity of the SCF, with three major classes of F-box proteins depending on the substrate interaction domain (WD40 repeats as βTrCP, leucine-rich repeats as SKP1 or SKP2, and other types such as Cyclin F). siRNA-based experiments showed the accumulation of SENP2 only in the cytosol of βTrCP knockdown cells (Figure 2I). Altogether, these results demonstrate that the SCF-E3 ligase Skp1-Cul-1-βTrCP mediates the degradation of SENP2 in the cytosol of cells.

The activity of SCF requires the conjugation of the ubiquitin-like polypeptide NEDD8 to Cul-1, suggesting the possibility that *Klebsiella* may trigger the loss of Cul-1 NEDDylation resulting in the lack of transfer of ubiquitin to SENP2. Typically, Cul-1 appears as a doublet at ∼85 and ∼90 kDa, with the higher molecular band representing the NEDDylated form of Cul-1 [22]. Immunoblotting experiments revealed the lack of Cul-1 NEDDylation in infected cells (Figure 2J). As a positive control, cells were pre-treated with H_2_O_2_ as this was shown to cause loss of Cul-1 NEDDylation (Figure 2J) [23].

DeNEDDylation of Cul-1 is normally catalysed by the COP9 signalosome (e.g., subunit 5 or CSN5/JAB1) [24]. We then examined whether repression of CSN5 expression may inhibit *Klebsiella*-induced Cul-1 deNEDDylation. As shown in Figure 2K, *Klebsiella* triggered Cul-1 deNEDDylation was significantly reduced in CSN5 knockdown cells. In CSN5 silenced cells *Klebsiella* did not reduce the K48-linked ubiquitylation of SENP2 (Figure 2L) and, as anticipated, there was a reduction in SENP2 levels in the cytosol of infected cells (Figure 2M). Finally, *Klebsiella* induced an increase in the global levels of SUMO1-conjugated proteins in CSN5 silenced cells (Figure 2N).

In summary, these results demonstrate that *K. pneumoniae* impairs the cytosolic degradation of SENP2 by affecting its K48-ubiquitylation through CSN5-mediated deNEDDylation of the Cul-1 subunit of the SCF E3 ligase Skp1-Cul-1-βTrCP.

### *K. pneumoniae* increases CSN5 levels through a EGFR-PI3K-AKT-ERK-GSK3β pathway

Transcription analysis showed that *K. pneumoniae* induced the expression of *csn5* in A549 cells (Figure 3A), and immunoblotting experiments demonstrated that Kp52145 increased the levels of CSN5 (Figure 3B). The regulation of CSN5 is poorly understood although there is evidence showing that CSN5 is a target of EGF receptor (EGFR) signalling [25]. Previous work from our laboratory revealed that *K. pneumoniae* activates the EGFR-phosphatidylinositol 3-OH kinase (PI3K)-AKT-ERK-GSK3β signalling pathway to limit inflammation [14]. We then investigated whether *Klebsiella*-induced expression of CSN5 is dependent on this signalling pathway. To test this hypothesis, we utilized pharmacologic inhibitors of EGFR (AG1478), PI3K (LY294002), AKT (AKT-X) or MEK (U0126) to block the pathway at four different levels. Immunoblotting revealed that Kp52145-triggered induction of CSN5 was reduced in cells pre-treated with the inhibitors (Figure 3C). Since we have shown that Kp52145 inactivates GSK3β by triggering the phosphorylation of serine 9 [17], we examined the role of inhibitory phosphorylation of GSK3β on CSN5 expression upon Kp52145 infection. Vectors encoding wild-type GSK3β (GSK3β-WT) or a constitutively active mutant (GSK3β−S9A) in which the serine 9 residue was changed to alanine were transiently transfected to A549 cells. As control, cells were transfected with the empty vector pcDNA3. *Klebsiella* did not induce CSN5 in cells transfected with the GSK3β-S9A vector (Figure 3C) demonstrating that the inhibitory phosphorylation of GSK3β is required for *Klebsiella* induction of CSN5. Next, we examined whether inhibition of this EGFR-controlled signalling pathway would abrogate *Klebsiella*-imposed reduction in SUMOylation. Indeed, pretreatment of the cells with the EGFR inhibitor AG1478 led to an overall increase in SUMO1 levels upon infection (Figure 3D). Because *Klebsiella* CPS is the bacterial factor triggering the activation of EGFR [14], we assessed whether *Klebsiella*-induced CSN5 up-regulation is dependent on the CPS. As anticipated, the *cps* mutant, strain 52145-Δ*wca*_*K2*,_ did not increase CSN5 levels in A549 cells (Figure 3E).

**Figure 3.**
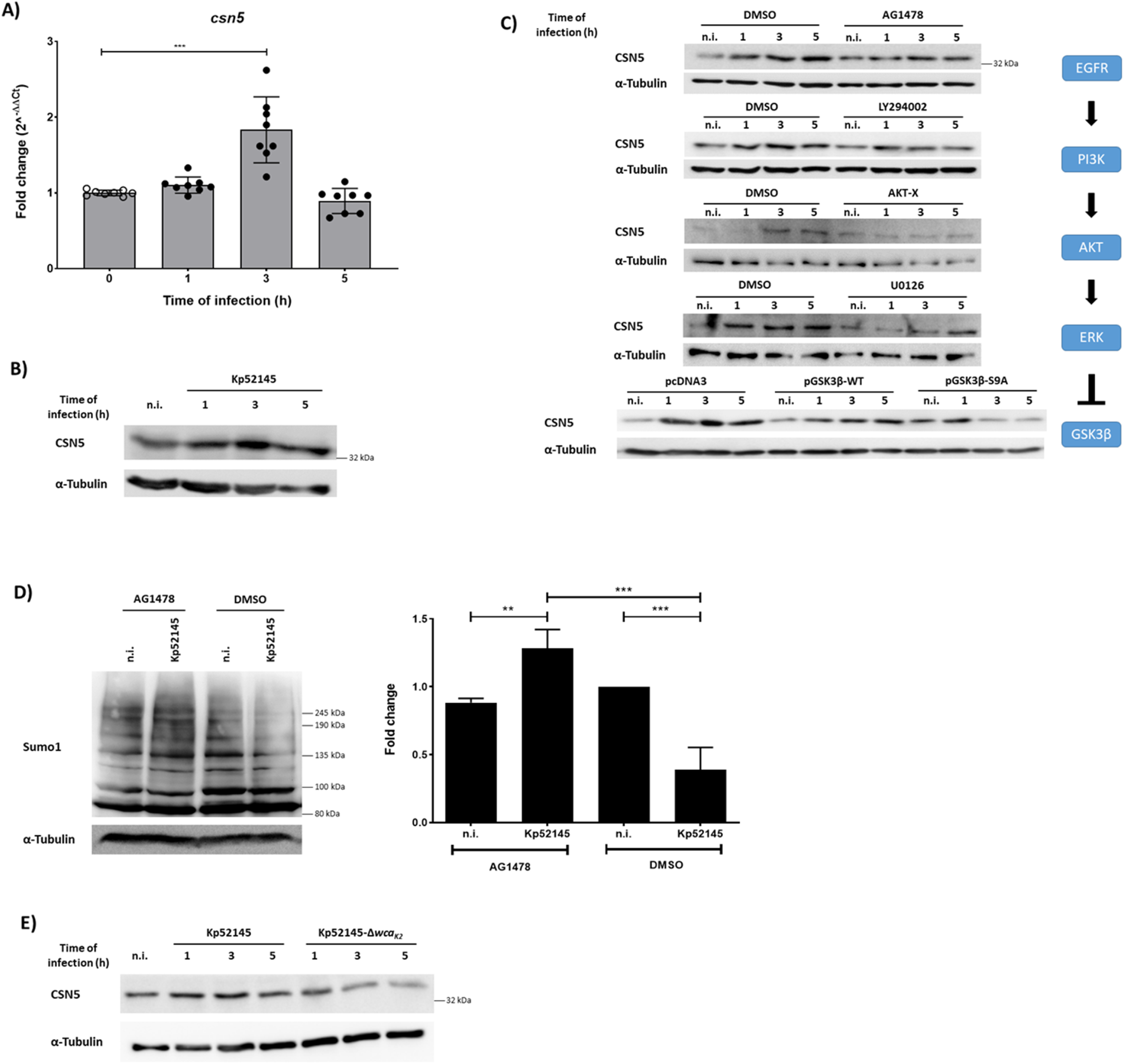
*K. pneumoniae* increases CSN5 levels in an EGFR-PI3K-AKT-ERK-GSK3β dependent manner. A. *csn5* mRNA levels, assessed by qPCR, in A549 cells left untreated (n.i.) or infected for up to 5 h with Kp52145. Values are presented as the mean ± SD of three independent experiments measured in duplicate. ***P ≤ 0.001 versus n.i. determined using one way-ANOVA with Bonferroni’s multiple comparisons test. B. Immunoblot analysis of CSN5 and tubulin levels in lysates of A549 cells infected with Kp52145 for the indicated times. C. Immunoblot analysis of CSN5 and tubulin levels in lysates of A549 cells infected with Kp52145 for the indicated times. Cells were pre-treated with AG1478 (5 μM), LY294002 (20 μM), AKT-X (30 μM), U0126 (10 μM) or DMSO (vehicle solution) for 2 h before infection where indicated. Where indicated cells were transfected with plasmids expressing GSK3β either wild-type or with a S9A mutation that renders the protein constitutively active or with pcDNA3, empty vector. The signalling pathway EGFR–PI3K–AKT–PAK4–ERK– GSK3β activated in *K. pneumoniae* infection is depicted to represent the several steps inhibited in this figure. D. Immunoblot analysis of SUMO1 levels in lysates of AG1478 (5 μM, 2 h before infection) or DMSO (vehicle solution)-treated A549 cells infected with Kp52145 for the indicated times. Membranes were reprobed for tubulin as a loading control. SUMO1 smears were quantified from three independent experiments using Image Studio Lite (Li-cor) and the modification normalised to α-Tubulin. The graph represents fold change at 5h of infection compared to DMSO non-infected control cells. **P ≤ 0.01 versus DMSO n.i. determined using one way-ANOVA with Bonferroni’s multiple comparisons test. E. Immunoblot analysis of CSN5 and tubulin levels in lysates of A549 cells infected with Kp52145 or 52145-Δ*wca*_*K2*_ for the indicated times. In all panels data are representative of at least three independent experiments.

Overall, these findings support the notion that *Klebsiella* CPS engages an EGFR-PI3K-AKT-ERK-GSK3β pathway to increase the levels of CSN5 to decrease the levels of SUMO-conjugated proteins.

### *K. pneumoniae* decreases SUMOylation in macrophages

Having demonstrated that *Klebsiella* reduces SUMOylation in epithelial cells, we investigated whether *Klebsiella* affects SUMO-conjugated proteins in macrophages. A wealth of evidence has established that macrophages play a crucial role in host defence against *K. pneumoniae* not only by clearing the bacteria but also by shaping the immune response against this pathogen [26,27]. MH-S mouse alveolar macrophages were infected with Kp52145 and the overall pattern of SUMO1-conjugated proteins was accessed by immunoblotting. Figure 4A shows that Kp52145 induced a decrease in the levels of SUMO1-conjugated proteins as well as a decrease in the levels of free SUMO1 (Figure 4B). Similar results were observed in infected THP-1 human macrophages (Figure 4C), indicating that the decrease in SUMOylation is not cell type dependent. Either UV-killed or heat-killed bacteria did not trigger a decrease in the level of SUMOylated proteins (Supplementary Figure S3) demonstrating that the reduction in the levels of SUMO1-conjugated proteins is mediated by live bacteria.

**Figure 4.**
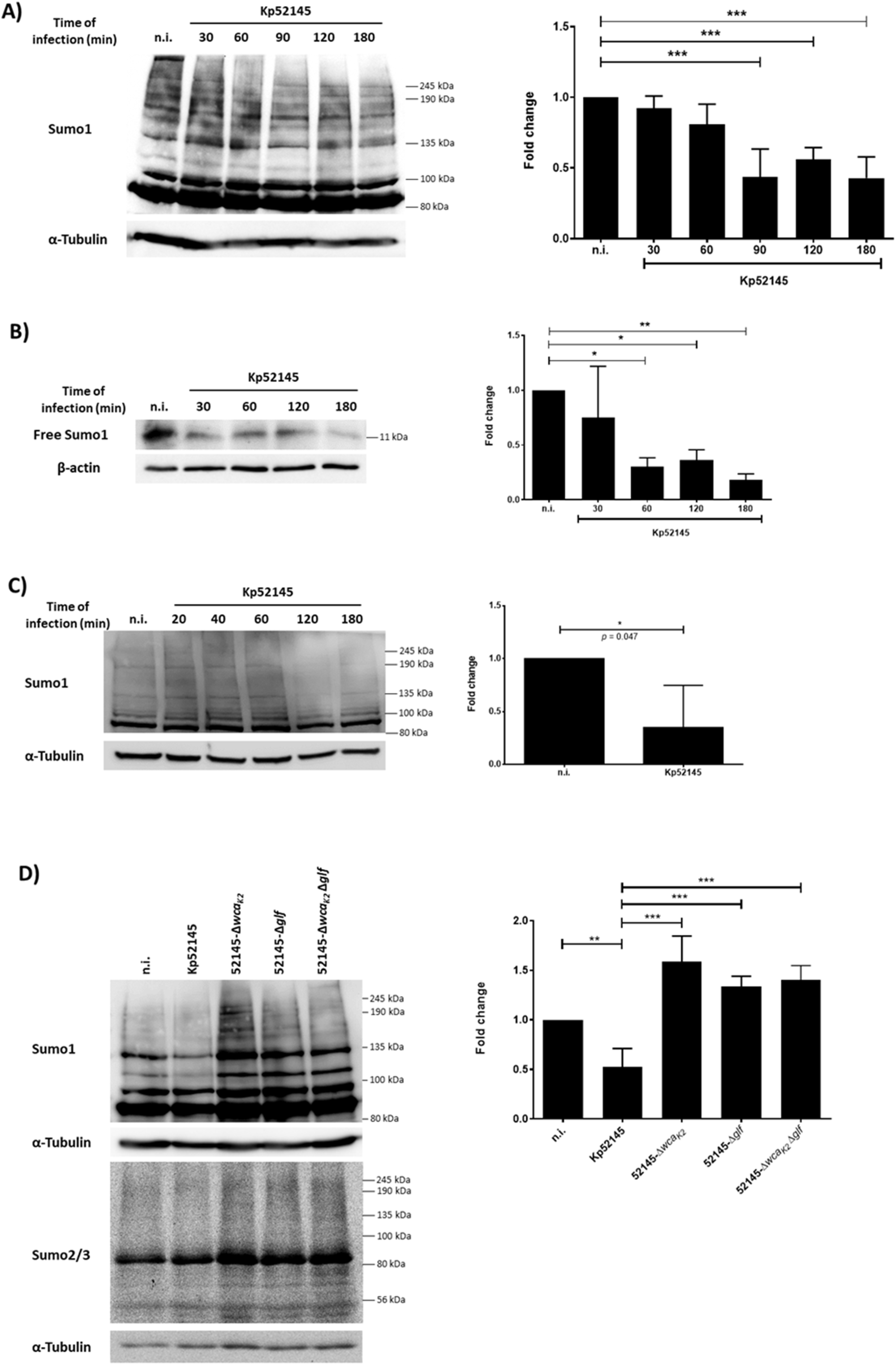
*K. pneumoniae* decreases SUMOylation in macrophages. A. Immunoblot analysis of SUMO1 and tubulin levels in lysates of mouse alveolar macrophages, MH-S cells infected with Kp52145 for the indicated time points. SUMO1 smears were quantified from three independent experiments using Image Studio Lite (Li-cor) and the modification normalised to α-Tubulin. The graph represents fold change compared to non-infected control cells at the indicated time points. **P ≤ 0.01; *P ≤ 0.05 versus n.i. determined using one way-ANOVA with Bonferroni’s multiple comparisons test. B. Immunoblot analysis of SUMO1 and tubulin levels in lysates of mouse alveolar macrophages, MH-S cells infected with Kp52145 for the indicated time points. The band corresponding to free SUMO1 at 11 kDa was quantified from three independent experiments using Image Studio Lite (Li-cor) and the modification normalised to α-Tubulin. The graph represents fold change compared to non-infected control cells at the indicated time points. *P ≤ 0.05 versus n.i. determined using one way-ANOVA with Bonferroni’s multiple comparisons test. C.Immunoblot analysis of SUMO1 and tubulin levels in lysates of human monocytic THP-1 cells infected with Kp52145 for the indicated time points. SUMO1 smears were quantified from three independent experiments using Image Studio Lite (Li-cor) and the modification normalised to α-Tubulin. The graph represents fold change at 180 min compared to non-infected control cells. *P = 0.0476 versus n.i. determined using two-tailed unpaired *t*-test. D.Immunoblot analysis of SUMO1 and tubulin levels in lysates of MH-S cells infected with Kp52145 or the indicated mutants for 1 h. SUMO1 smears were quantified from three independent experiments using Image Studio Lite (Li-cor) and the modification normalised to α-Tubulin. The graph represents fold change compared to non-infected control cells. *P ≤ 0.05 versus n.i. determined using one way-ANOVA with Bonferroni’s multiple comparisons test. n.i. – non-infected control In all panels data are representative of at least three independent experiments.

Previous work from this laboratory demonstrated the importance of the *K. pneumoniae* polysaccharides, CPS and lipopolysaccharide (LPS), in the interplay between the pathogen and macrophages [28]. To examine whether CPS and LPS mediate the *Klebsiella*-triggered decrease in SUMOylation, we assessed the global pattern of SUMO1-conjugated proteins in macrophages infected with the CPS (52145-Δ*wca*_*K2*_), LPS O-polysaccharide (52145-Δ*glf*) mutants and the double mutant lacking both polysaccharides (52145-Δ*wca*_*K2*_ Δ*glf*). Immunoblotting revealed an increase in the levels of SUMO-conjugated proteins in macrophages infected with the CPS and LPS O-polysaccharide mutants (Figure 4D), indicating that both polysaccharides have a role in the *Klebsiella*-triggered decrease in SUMOylated proteins. Controls confirmed there were no differences in the levels of surface attached capsule produced by the wild-type and the LPS O-polysaccharide mutant (174.4 and 174.8 μg glucuronic acid.10^−9^ CFU, respectively).

### *K. pneumoniae* induced decrease in SUMOylation in macrophages is mediated by TLR4-dependent type-I interferon

Like in epithelial cells, Ubc9 levels were not decreased in *Klebsiella*-infected macrophages (Supplementary Figure S4A). Cell fractionation experiments did not reveal increased levels of SENP2 in the cytosol (Supplementary Figure S4B), and NEDDylation of Cul-1 was not impaired in infected macrophages (Supplementary Figure S4C). Collectively, these observations indicated that *Klebsiella* may employ a different strategy in macrophages to decrease the levels of SUMO1-conjugated proteins.

The fact that the CPS and LPS O-polysaccharide mediated the *Klebsiella*-triggered decrease in SUMOylated proteins and that both polysaccharides are recognised by TLR4 [13,29] led us to investigate whether *Klebsiella* targets TLR4 signalling to decrease SUMOylation. We observed an increase in SUMOylated proteins in infected *TLR4*^-/-^ iBMDMs (Immortalised bone marrow-derived macrophages) (Figure 5A). TLR4 signals via adaptors MyD88 and TRIF, with TRAM acting as a sorting adaptor controlling TRIF localisation [30]. To learn which TLR4 adaptor(s) mediate(s) *Klebsiella*-induced decrease in global SUMOylation, *MyD88*^-/-^ and *TRAM/TRIF*^-/-^ iBMDMs were infected. We observed no differences in the global levels of SUMO1-conjugated proteins between wild-type and *MyD88*^-/-^ iBMDMs (Figure 5B). In contrast, there was an increase in the levels of SUMOylated proteins in *TRAM/TRIF*^-/-^ iBMDMs (Figure 5C), indicating that *Klebsiella*-triggered decrease in SUMOylation is dependent on TLR4-TRAM-TRIF signalling.

**Figure 5.**
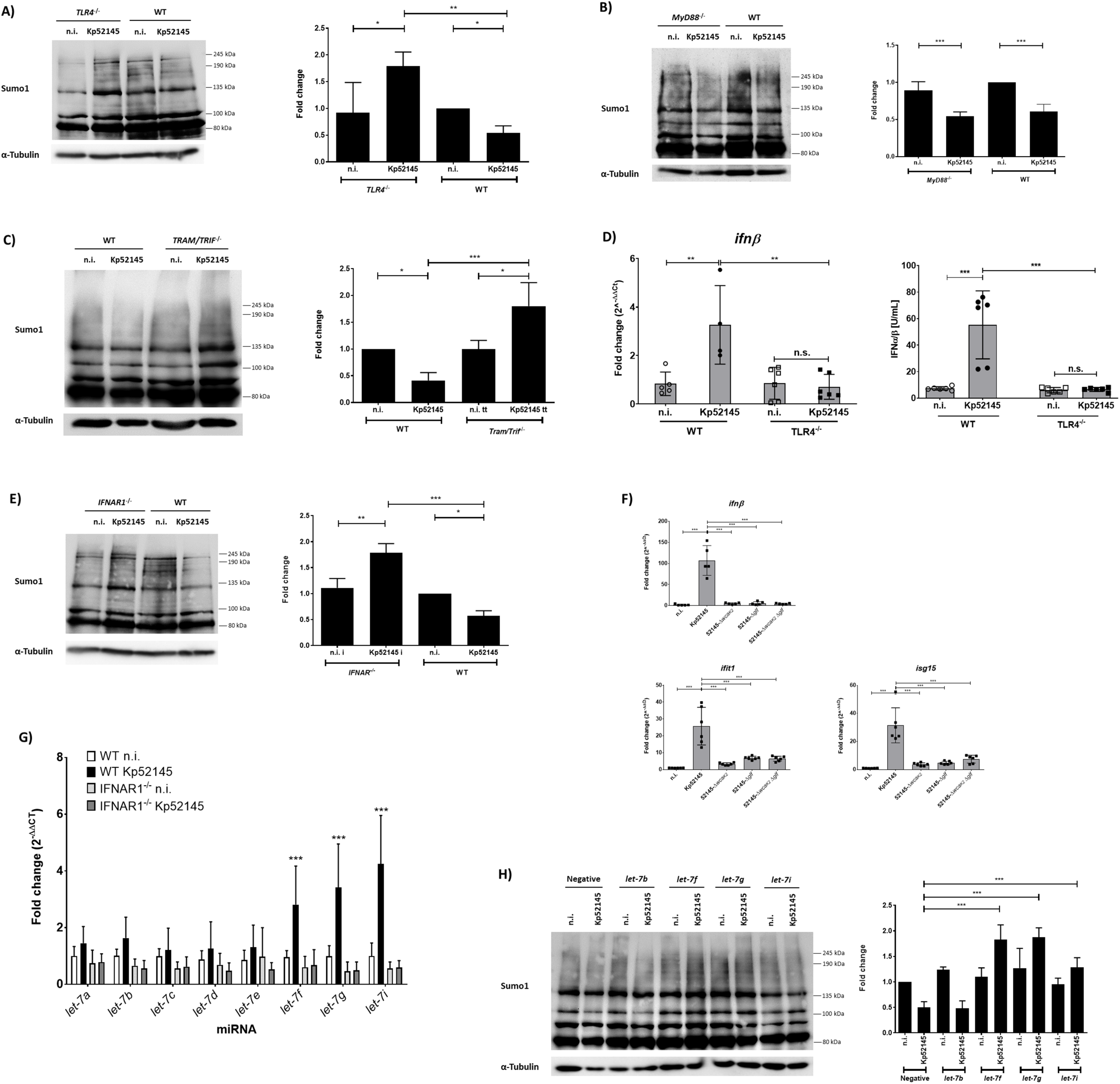
*K. pneumoniae* SUMOylation decrease in macrophages is mediated by TLR4-dependent type-I interferon and *let-7* miRNAs. A. Immunoblot analysis of SUMO1 and tubulin levels in lysates of immortalised bone marrow-derived macrophages, iBMDM cells derived from C57BL/6 mice (WT) or *Tlr4* knockout mice (*TLR4*^-/-^) infected with Kp52145 for the indicated time points. SUMO1 smears were quantified from three independent experiments using Image Studio Lite (Li-cor) and the modification normalised to α-Tubulin. The graph represents fold change compared to WT iBMDM non-infected control cells. ***P ≤ 0.001; **P ≤ 0.01; *P ≤ 0.05 versus WT n.i. determined using one way-ANOVA with Bonferroni’s multiple comparisons test. B. Immunoblot analysis of SUMO1 and tubulin levels in lysates of iBMDM cells derived from C57BL/6 mice (WT) or MyD88 knockout mice (*MyD88*^-/-^) infected with Kp52145 for the indicated time points. SUMO1 smears were quantified from three independent experiments using Image Studio Lite (Li-cor) and the modification normalised to α-Tubulin. The graph represents fold change compared to WT iBMDM non-infected control cells. **P ≤ 0.01; *P ≤ 0.05 versus WT n.i. determined using one way-ANOVA with Bonferroni’s multiple comparisons test. C. Immunoblot analysis of SUMO1 and tubulin levels in lysates of iBMDM cells derived from C57BL/6 mice (WT) or *Tram/Trif* double knockout mice (*TRAM/TRIF*^-/-^) infected with Kp52145 for the indicated time points. SUMO1 smears were quantified from three independent experiments using Image Studio Lite (Li-cor) and the modification normalised to α-Tubulin. The graph represents fold change compared to WT iBMDM non-infected control cells. ***P ≤ 0.001; *P ≤ 0.05 versus WT n.i. determined using one way-ANOVA with Bonferroni’s multiple comparisons test. D. Type I-IFN levels determined in the supernatants of iBMDM WT or TLR4^-/-^ cells left untreated (n.i.) or infected for 3 h with Kp52145. The reporter cell line B16-Blue™ IFN-α/β was used for the quantification of levels of SEAP produced upon stimulation of the supernatants with the detection medium QUANTI-Blue™ and presented as IFNα/β U mL^-1^. *IFNβ* mRNA levels were assessed by qPCR and are presented as fold change against untreated WT cell levels. Values are presented as the mean ± SD of three independent experiments measured in duplicate. ***P ≤ 0.001; *P ≤ 0.05; versus n.i. determined using one way-ANOVA with Tukey’s multiple comparisons test. n.s. – non significant E. Immunoblot analysis of SUMO1 and tubulin levels in lysates of BMDM cells derived from C57BL/6 mice (WT) or IFNAR1 knockout mice (*IFNAR1*^-/-^) infected with Kp52145 for the indicated time points or left untreated (n.i.). SUMO1 smears were quantified from three independent experiments using Image Studio Lite (Li-cor) and the modification normalised to α-Tubulin. The graph represents fold change compared to WT BMDM non-infected control cells. **P ≤ 0.01; * ≤ 0.05 versus WT n.i. determined using one way-ANOVA with Bonferroni’s multiple comparisons test. F.*ifnβ, ifit1* and *isg15* mRNA levels, assessed by qPCR, in MH-S cells left untreated (n.i.) or infected for 3 h with the *K. pneumoniae* strains indicated. Values are presented as the mean ± SD of three independent experiments measured in duplicate. ***P ≤ 0.001; *P ≤ 0.05; versus n.i. determined using one way-ANOVA with Tukey’s multiple comparisons test. G.*let-7* mRNA levels, assessed by qPCR, in BMDM cells derived from C57BL/6 mice (WT) or IFNAR1 knockout mice (*IFNAR1*^-/-^) infected with Kp52145 for 1 h or left untreated (n.i.). Values are presented as the mean ± SD of three independent experiments measured in duplicate. ***P ≤ 0.001; *P ≤ 0.05; versus iBMDM WT n.i. determined using two way-ANOVA with Holm-Sidak’s multiple comparisons test. H.Immunoblot analysis of SUMO1 and tubulin levels in lysates of *let-7* miRNA antagomir transfected MH-S cells infected with Kp52145 for 1 h or left untreated (n.i.). Negative – *Caenorhabditis elegans* control sequence with minimal sequence identity in mouse cells. SUMO1 smears were quantified from three independent experiments using Image Studio Lite (Li-cor) and the modification normalised to α-Tubulin. The graph represents fold change compared to Negative-transfected non-infected control cells. ***P ≤ 0.001; **P ≤ 0.01; versus Negative n.i. determined using one way-ANOVA with Bonferroni’s multiple comparisons test. In panels A, B, C, E and H, data are representative of at least three independent experiments.

The TLR4-TRAM-TRIF signalling pathway governs type-I IFN production [31], making it relevant to investigate whether the *Klebsiella-*induced decrease in SUMOylation is mediated by type-I IFN. Further confirming recent findings of our laboratory [32], Kp52145 induced IFNβ, *ifnβ* and interferon-stimulated genes (ISGs) in wild-type iBMDMs but not in *TLR4*^-/-^ (Figure 5D and Supplementary Figure S5A). To demonstrate the contribution of type-I IFN to the *Klebsiella*-triggered decrease in levels of SUMO-conjugated proteins, we infected BMDMs obtained from *IFNAR1*^-/-^ mice. Kp52145 increased the levels of SUMOylated proteins in *IFNAR1*^-/-^ BMDMs (Figure 5E), demonstrating that the *Klebsiella*-induced decrease in SUMOylation is indeed dependent on type-I IFN signalling. We then investigated whether a reduced induction of type-I IFNs by the CPS and LPS O-polysaccharide mutants may explain the high levels of SUMO1-conjugated proteins in infected macrophages. Indeed, the three mutants induced less *ifnβ* and ISGs *ifit1* and *isg15* expression than the wild type (Figure 5F). Furthermore, addition of type-I IFN to macrophages infected with the double mutant lacking CPS and LPS O-polysaccharide rescued the decrease in the levels of SUMOylated proteins (Supplementary Figure S5B). Altogether, these findings provide compelling evidence to the notion that TLR4-TRAM-TRIF induced type-I IFN via IFNAR1-controlled signalling mediates *Klebsiella*-triggered decrease in the levels of SUMOylation.

### *let-7* miRNAs mediate type-I IFN decrease in SUMOylation

IFNs regulate the expression of miRNAs from the *let-7* family [33]. miRNAs downregulate mRNA stability and protein synthesis through complementary elements in the 3′-untranslated regions (UTRs) of their target mRNAs. Sahin and coworkers reported that type-I IFNs control SUMO expression and validated *let-7* target sequences in 3’UTRs of human and mouse *sumo1, sumo2* and *sumo3* transcripts [34], indicating an evolutionary conserved regulation of SUMO by *let-7* family members. This evidence prompted us to investigate whether the *Klebsiella*-induced decrease in the overall levels of SUMO-conjugated proteins by type-I IFNs is through *let-7* miRNAs. We reasoned that Kp52145 should upregulate the expression of *let-7* miRNA member(s) only in wild-type but not in *IFNAR1*^-/-^ BMDMs. Figure 5G shows that *Klebsiella* upregulated the expression of *let-7f, let-7g* and *let-7i* in wild-type BMDMs. These miRNAs were not upregulated in *Klebsiella* infected *IFNAR1*^-/-^ BMDMs (Figure 5G), leading us to study whether these *let-7* family members mediate type-I IFN-controlled decrease in SUMOylation in *Klebsiella*-infected macrophages. To validate this hypothesis, antagomirs for these *let-7* family members were transfected into macrophages and the global pattern of SUMOylation determined by immunoblotting. As controls, we also tested a non-targeting control antagomir and a *let-7b* antagomir since this miRNA was not significantly upregulated by *Klebsiella* in wild-type BMDMs (Figure 5G). In contrast to cells transfected with a non-targeting control, inhibition of *let-7f, let-7g* and *let-7i* led to a global upregulation of SUMOylation in infected cells (Figure 5H). The effect was less prominent in *let-7i* inhibited cells than in *let-7f* and *let-7g* ones (Figure 5H). Collectively, these results reveal that *Klebsiella* dependent type-I IFN-decrease in SUMOylation involves *let-7* miRNAs.

### SUMOylation increases host defences against *K. pneumoniae*

The sophisticated strategies employed by *Klebsiella* to decrease the levels of SUMO-conjugated proteins in epithelial cells and macrophages strongly suggested that SUMOylation should play an important role in host defence against *Klebsiella* infections. Therefore, we sought to determine whether alterations in the levels of SUMO1-conjugated proteins might influence *Klebsiella*-cell cross talk. By using a SUMO1 expression plasmid, a common method to upregulate the SUMO status of the host proteins [5,6], we transiently upregulated SUMOylation (Supplementary Figure S6A and S6B). We and others have established that dampening inflammation is a hallmark of *Klebsiella* infection biology *in vitro* and *in vivo* [12,14,15,35,36]. Therefore, we examined the levels of IL-8 and TNFα released by infected epithelial cells and macrophages, respectively. IL-8 and TNFα are well-established inflammatory read-outs upon infection. In SUMO1-overexpressing A549 cells, we observed a significant increase in IL-8 in the supernatants of infected cells (Figure 6A). Furthermore, knockdown of SENP2 also leads to increased release of IL-8, strengthening the role of this protein in controlling SUMOylation levels and inflammation (Figure 6B). SUMO1-overexpressing macrophages also produced more TNFα upon infection (Figure 6C). Interestingly, macrophages transfected with *let-7f* and *let-7g* antagomirs also released more TNFα upon infection (Figure 6D) further sustaining the connection between increased SUMOylation and inflammation in macrophages. Collectively, these findings suggest that *Klebsiella*-triggered decrease in SUMOylation in epithelial cells and macrophages is a pathogen strategy to limit inflammation.

**Figure 6.**
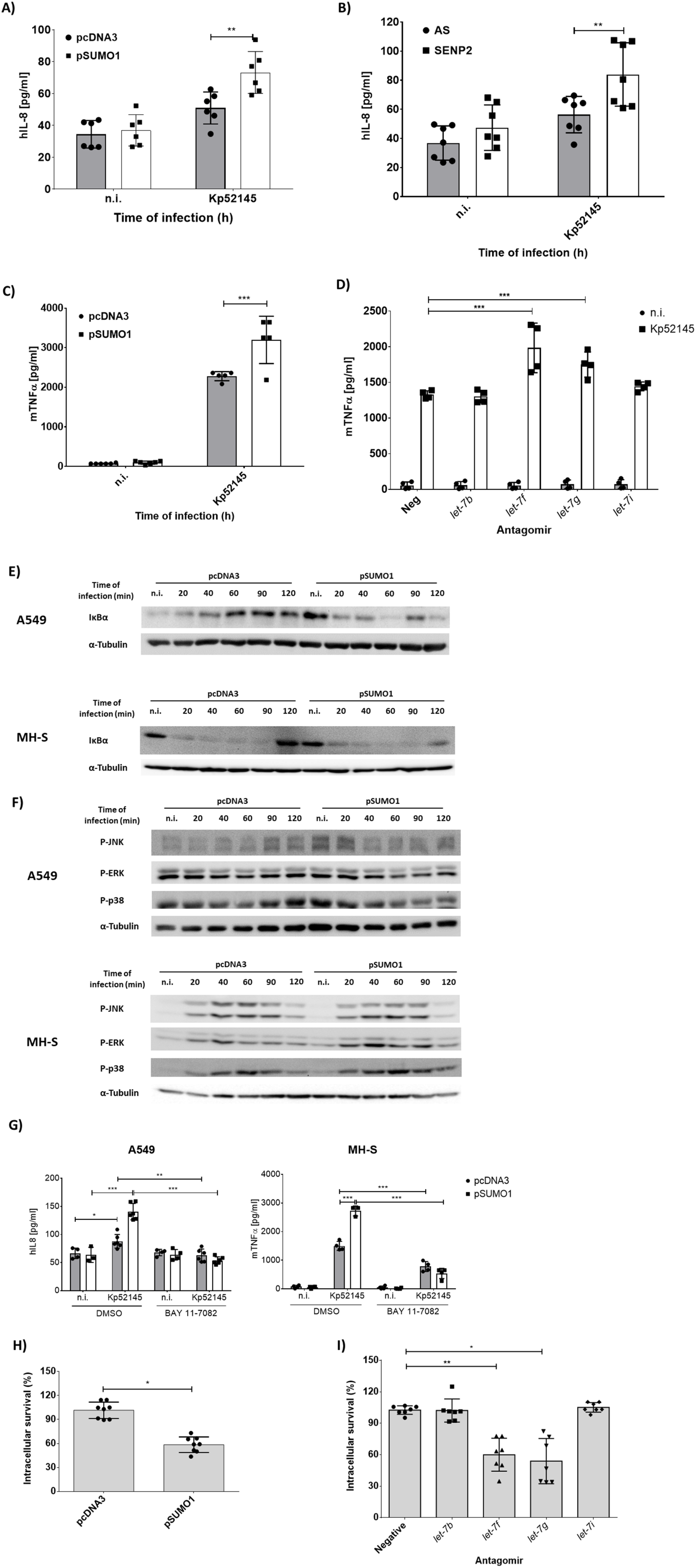
SUMOylation increases host defences against *K. pneumoniae*. A. ELISA of IL-8 secreted by A549 cells transfected with either empty control plasmid (pcDNA3) or SUMO1-HA tagged plasmid (pSUMO1), which were left untreated (n.i.) or infected with Kp52145 for 3 h of contact after which the medium was replaced with medium containing gentamicin (100 µg mL^-1^) to kill extracellular bacteria and incubated for a further 4 hours. Values are presented as the mean ± SD of three independent experiments measured in duplicate. ***P ≤ 0.001; *P ≤ 0.05; versus pcDNA3 determined using two way-ANOVA with Holm-Sidak’s multiple comparisons test. B. ELISA of IL-8 secreted by control (AS) or Senp2 siRNA-transfected A549, which were left untreated (n.i.) or infected with Kp52145 for 3 h of contact after which the medium was replaced with medium containing gentamicin (100 µg mL^-1^) to kill extracellular bacteria and incubated for a further 4 hours. Values are presented as the mean ± SD of three independent experiments measured in duplicate. ***P ≤ 0.001; *P ≤ 0.05; versus AS determined using two way-ANOVA with Holm-Sidak’s multiple comparisons test. C.ELISA of TNFα secreted by empty control (pcDNA3) or SUMO1-HA plasmid-transfected MH-S, which were left untreated (n.i.) or infected with Kp52145 for 1 h of contact after which the medium was replaced with medium containing gentamicin (100 µg mL^-1^) to kill extracellular bacteria and incubated for a further 4 hours. Values are presented as the mean ± SD of three independent experiments measured in duplicate. ***P ≤ 0.001; *P ≤ 0.05; versus pcDNA3 determined using two way-ANOVA with Holm-Sidak’s multiple comparisons test. D.ELISA of TNFα secreted by antagomir-transfected MH-S, which were left untreated (n.i.) or infected with Kp52145 for 1 h of contact after which the medium was replaced with medium containing gentamicin (100 µg mL^-1^) to kill extracellular bacteria and incubated for a further 4 hours. Values are presented as the mean ± SD of three independent experiments measured in duplicate. ***P ≤ 0.001; *P ≤ 0.05; versus Negative Kp52145 determined using two way-ANOVA with Holm-Sidak’s multiple comparisons test. E.Immunoblot analysis of IκBα and tubulin levels in lysates of empty (pcDNA3) or SUMO1-HA (pSUMO1) plasmid transfected A549 or MH-S cells, infected with Kp52145 for the indicated time points. F.Immunoblot analysis of JNK (P-JNK), ERK (P-ERK) and p38 (P-p38) phosphorylation levels in lysates of empty (pcDNA3) or SUMO1-HA (pSUMO1) plasmid transfected A549 or MH-S cells, infected with Kp52145 for the indicated time points. Tubulin immunoblotting was used as the loading control. n.i. – non-infected control G.ELISA of IL-8 or TNFα secreted by empty control (pcDNA3) or SUMO1-HA plasmid-transfected A549 or MH-S respectively, which were treated with the NF-κB inhibitor BAY 11-7082 (5 μM, 2 h before infection) or DMSO (vehicle solution). Cells were left untreated (n.i.) or infected with Kp52145 for 1 h (MH-S) or 3 h (A549) of contact after which the medium was replaced with medium containing gentamicin (100 µg mL^-1^) to kill extracellular bacteria and incubated for a further 4 hours. Values are presented as the mean ± SD of three independent experiments measured in duplicate. ***P ≤ 0.001; *P ≤ 0.05; determined using two way-ANOVA with Holm-Sidak’s multiple comparisons test. H.Percentage of intracellular survival in MH-S cells transfected with either empty control plasmid (pcDNA3) or SUMO1-HA tagged plasmid (pSUMO1). Cells were infected with Kp52145 for 30 min (MOI 100:1), wells were washed and incubated with medium containing gentamicin (100 μg mL^-1^) for 2.5 h. Intracellular bacteria were quantified by lysis, serial dilution and viable counting on LB agar plates. Percentage of intracellular survival was determined by dividing the number of colony forming units (CFU) obtained at 3 h of infection over the number obtained at 1 h and considering pcDNA3 as 100 %. Values are presented as the mean ± SD of three independent experiments measured in duplicate. ***P ≤ 0.001; *P ≤ 0.05; versus pcDNA3 determined using unpaired t-test with correction for Holm-Sidak’s multiple comparisons test. I.Percentage of intracellular survival in antagomir transfected MH-S cells. Cells were infected with Kp52145 for 30 min (MOI 100:1), wells were washed and incubated with medium containing gentamicin (100 μg mL^-1^) for 2.5 h. Intracellular bacteria were quantified by lysis, serial dilution and viable counting on LB agar plates. Percentage of intracellular survival was determined by dividing the number of colony forming units (CFU) obtained at 3 h of infection over the number obtained at 1 h and considering the negative control as 100 %. Values are presented as the mean ± SD of three independent experiments measured in duplicate. **P ≤ 0.01; *P ≤ 0.05; versus negative control determined using one way-ANOVA with Holm-Sidak’s multiple comparisons test. In panels E and F, data are representative of at least three independent experiments.

To further investigate this notion, we examined the activation of NF-κB and MAP kinases by *Klebsiella*. NF-κB and MAP kinases govern the expression of inflammatory markers upon *Klebsiella* infection [13,15]. In the canonical NF-κB activation pathway, nuclear translocation of NF-κB is preceded by phosphorylation and subsequent degradation of IκBα. IκBα levels were reduced in SUMO1-overexpressing infected epithelial cells and macrophages (Figure 6E). No significant differences in the phosphorylation of MAPKs were observed (Figure 6F), indicating that the activation status of MAPKs upon *Klebsiella* infection is not affected by SUMOylation. The inhibition of the canonical NF-κB pathway with BAY 11-7082 led to a marked decrease of cytokine release in SUMO1-overexpressing cells (Figure 6G), connecting the increased release of inflammatory mediators in SUMO1-overexpressing cells with the activation of NF-κB.

Recently, we have demonstrated that *K. pneumoniae* survives inside macrophages residing in the *Klebsiella*-containing vacuole [37]. To learn whether SUMOylation plays any role in *Klebsiella* intracellular survival, SUMO1-overexpressing macrophages were infected, and the number of intracellular bacteria was determined 3h post infection. Figure 6H shows that the number of intracellular bacteria was significantly reduced in cells overexpressing SUMO. *Klebsiella* attachment was not affected in SUMO1 overexpressing cells (Supplementary Figure S7C), indicating that an increase in SUMOylation is detrimental for *K. pneumoniae* intracellular survival. Further sustaining this notion, we observed a significant decrease in bacterial numbers in macrophages transfected with *let-7f* and *let-7g* antagomirs (Figure 6I). *Klebsiella* attachment was not affected in antagomir transfected cells (Supplementary Figure S6D), demonstrating the importance of *Klebsiella*-induced *let-7* family members for intracellular survival.

Overall, these data provide evidence that *Klebsiella*-induced decrease in SUMOylation promotes bacterial infection.

## DISCUSSION

SUMOylation is a conserved process in the eukaryotic kingdom. In humans, thousands of SUMOylated proteins have been identified, and they are involved in transcription regulation, stress responses and cell intrinsic immunity among many other biological processes [1]. Consistent with the essential role of SUMOylation in the host cell, it has been shown that pathogens can interfere with this PTM to promote their own survival and replication. The vast majority of evidence refers to how viruses target SUMOylation [4], whereas little is known concerning the interplay between bacteria and this PTM. In this work, we demonstrate that the human pathogen *K. pneumoniae* induces a decrease in SUMOylation in epithelial cells and macrophages for intracellular survival and to limit the activation of inflammatory responses. Mechanistically, our findings uncover hitherto unknown strategies exploited by a pathogen to interfere with SUMOylation (Figure 7). In epithelial cells, *K. pneumoniae* hijacks the SENP2 deSUMOylase whereas in macrophages *Klebsiella* exploits type-I IFN-induced *let-7* miRNA to alter SUMOylation.

**Graphical abstract / Figure 7.**
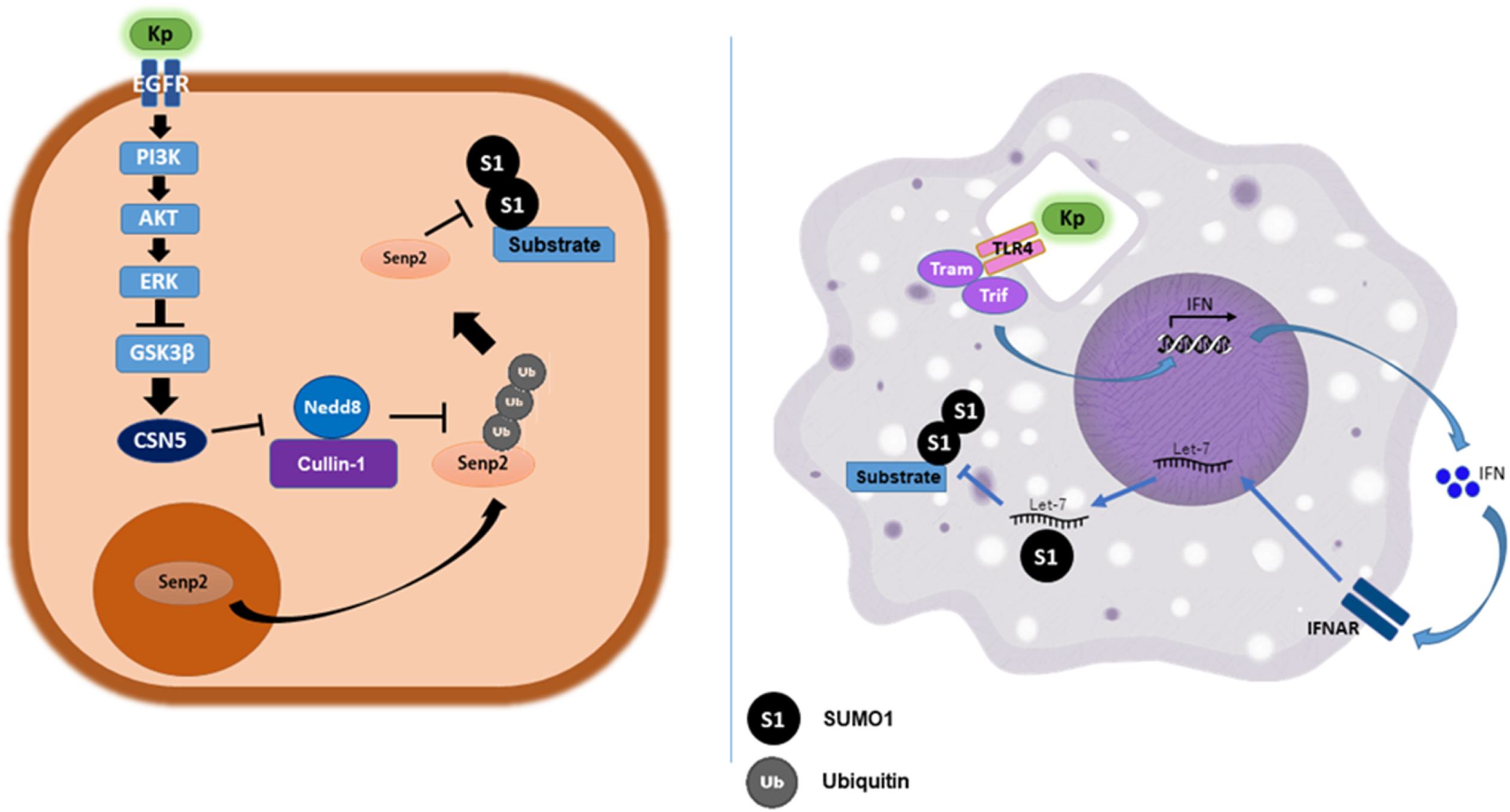
*Klebsiella pneumoniae* ablates SUMOylation via distinct pathways in epithelial cells or macrophages. Working model of *K. pneumoniae* strategies to decrease SUMOylation in epithelial cells and macrophages. In epithelial cells, Kp52145 activates the signalling pathway EGFR-PI3K-AKT-ERK-GSK3β to increase the COP9 signalosome component CSN5 which inhibits NEDDylation of Cullin-1 and prevents proteasomal degradation of SENP2. SENP2 then accumulates in the cytosol preventing SUMOylation. In macrophages, Kp52145 via TLR4-TRAM-TRIF induces the production of type-I interferon, which signals through the IFNAR1 receptor. Type-I interferon stimulates transcription of the miRNA *let-7* which prevents SUMOylation. Both strategies lead to increased intracellular survival and subversion of host responses.

Our results in epithelial cells are reminiscent of those reported for the intracellular pathogens *L. monocytogenes, S. flexneri* and *S. typhimurium* that, likewise *Klebsiella*, decrease the levels of SUMO-conjugated proteins. Mechanistically, these pathogens induce the degradation of the E1 enzyme SAE2 and E2 enzyme Ubc9 to decrease SUMOylation [5– 8]. However, and to the best of our knowledge, *Klebsiella*, a pathogen that does not invade epithelial cells, is the first among viral and bacterial pathogens exploiting a host deSUMOylase to reduce SUMOylation. Given the global decrease in levels of SUMO1-conjugated proteins triggered by *Klebsiella*, it was puzzling to identify the nuclear associated deSUMOylase SENP2 as the deSUMOylase targeted by the pathogen. Several lines of evidence sustained that *Klebsiella* prevents the ubiquitin proteasome dependent degradation of SENP2 to increase its levels in the cytosol to decrease SUMOylation. Our results conclusively demonstrate that *Klebsiella* targets NEDDylation, another PTM, to abrogate the K48-linked ubiquitylation of SENP2. To date, only commensal non-pathogenic bacteria have been shown to trigger the loss of NEDDylated Cul-1 in epithelial cells via generation of reactive oxygen species which affect the function of the E2 enzyme Ubc12 [23], whereas the CHBP and Cif type 3 secretion effectors from *Burkholderia pseudomallei* and enteropathogenic *Escherichia coli*, respectively, trigger the deamidation of NEDD8 to abolish the ubiquitin ligase activity of the E3 ubiquitin ligase complex [37]. In contrast, *Klebsiella*-induced lack of NEDDylation of Cul-1 was dependent on the CSN5 deNEDDylase. Our findings demonstrate that *Klebsiella* induces CSN5 in an EGFR-PI3K-AKT-ERK-GSK3β pathway dependent manner. Interestingly, *Klebsiella* activates this pathway to blunt inflammation by inducting the expression of the deubiquitylase CYLD to limit the ubiquitylation of TRAF6 [15], placing this EGFR-governed pathway at the fulcrum of *Klebsiella* strategies to control PTMs to promote infection.

The fact that NEDDylation of Cul-1 is essential for the function of the ubiquitin ligase complex E3-SCF^βTrCP^ that controls the levels of proteins involved in host-pathogen interactions such as p53, β-catenin, and IκBα among many others [38], warrants future studies to investigate the levels of proteins controlled by regulated proteolysis in *Klebsiella* infected epithelial cells. Interestingly, recent proteomic analysis have revealed the NEDDylation of several hundreds of proteins [39]. It is beyond the scope of this work to assess the impact of *K. pneumoniae* on the NEDDylation of proteins other than Cul-1. However, and based on the results of this work, we hypothesise that *Klebsiella* may decrease global NEDDylation which may also contribute to the panoply of strategies developed by *Klebsiella* to manipulate host cell biology.

A common theme of the interplay between *K. pneumoniae* and epithelial cells emerges from this work and previous studies from our laboratory [14,15]. Here, we have shown that *Klebsiella* hijacks two proteins, SENP2 and CSN5, crucial to allow reversible conjugation of SUMO1 and NEDD8 to target proteins, respectively. Previously, we demonstrated that *Klebsiella* blocks the activation of NF-κB and MAPK signalling pathways via the deubiquitinase CYLD and the phosphatase MKP-1, respectively by preventing the ubiquitylation and phosphorylation of key signalling molecules [15]. Cells activate CYLD and MKP1 to return to homeostasis after inflammation to protect the host from an overwhelming inflammatory response [40,41]. Therefore, the picture that is taking shape is that a signature of *K. pneumoniae* infection biology is interfering with PTMs by hijacking host proteins used by the cells to return to homeostasis. This strategy is radically different to that employed by other pathogens such as *Listeria, Salmonella, Shigella* or *Mycobacterium* who deliver bacterial proteins into host cells to modulate PTMs.

Another novel finding of this work is that *K. pneumoniae* interfered with SUMOylation in macrophages. Despite the crucial role played by macrophages in host defence and tissue homeostasis, there is limited evidence of pathogen-mediated alteration of PTMs in this cell type other than phosphorylation and ubiquitylation of proteins involved in the control of inflammatory responses. In this work, we have uncovered that the *Klebsiella*-triggered decrease of SUMOylation was dependent on TLR4-TRAM-TRIF induced type-I IFN. Whereas the role of SUMOylation governing IFN production during viral infection is well appreciated [42], we are not aware of any pathogen exploiting type-I IFN to decrease the levels of SUMO-conjugated proteins. In fact, the role of type-I IFNs in bacterial infections is still poorly understood [31], playing an adverse role in certain bacterial infections, while in others type-I IFNs are critical for host defence. In any case, our findings further expand the regulatory functions of type-I IFNs beyond the control of innate and adaptive immune responses [43]. Our results support the concept that the decrease in SUMOylation dependent on *Klebsiella* induced type-I IFN involves the upregulation of *let-7* miRNAs. This result is in line with previous studies showing post-transcriptional regulation of SUMO levels by *let-7* miRNA family members. It is interesting to note that depletion of Ubc9 by *S. typhimurium* relies on upregulation of the small noncoding RNAs *miR-30c* and *miR-30e* [6] whereas Epstein-Barr virus encoded miRNAs target several members of the SUMO interactome [44]. It is therefore tempting to speculate that post-transcriptional regulation of components of the SUMOylation cascade might be a general strategy shared by pathogens to decrease SUMOylation. Some miRNAs, including *miR-146, miR-155, miR-125, let-7* and *miR-21*, are commonly affected during bacterial infection to modulate host inflammatory responses [45]. Interestingly, evidence indicates that downregulation of *let-7* is a conserved response shared by different pathogens to sustain inflammation [45,46]. In stark contrast, here, we have shown that *K. pneumoniae* upregulates the expression of *let-7* miRNA family members in a type-I IFN-dependent manner. Moreover, abrogation of these miRNAs by transfecting specific antagomirs resulted in an increase of inflammatory responses and decreased intracellular survival in macrophages, supporting the notion that *K. pneumoniae* exploits *let-7* miRNA to manipulate host pathways. Future studies are warranted to develop a more comprehensive understanding of the roles of miRNAs in the *Klebsiella*-host interplay.

We were keen to identify the *K. pneumoniae* factor(s) mediating the *Klebsiella*-triggered decrease in SUMOylation. Our results revealed the crucial role played by *Klebsiella* polysaccharides, the CPS and the LPS O-polysaccharide, to decrease the levels of SUMO-conjugated proteins. The CPS and the LPS are perhaps two of the best-characterised virulence factors of *K. pneumoniae*; the findings of our work expand the roles of these polysaccharides to counteract host defence responses. Importantly, these polysaccharides are required for *K. pneumoniae* survival in mice (pneumonia model) [12,47,48], underlining the importance of SUMOylation reduction as a *Klebsiella* virulence trait since this process is abrogated in these mutant strains. In epithelial cells, CPS limits SUMOylation by preventing bacterial internalisation, and increasing the levels of the deNEDDylase CSN5. Notably, changes in the global pattern of SUMO-conjugated proteins in *Listeria* and *Shigella* infection affect internalisation of bacteria [5,49]; and, in *Klebsiella* infection, internalisation of the *cps* mutant is associated with an increase in SUMOylation in epithelial cells. The fact that *Klebsiella, Listeria* and *Shigella* decrease SUMOylation over time [5,48] suggests that the proteins SUMOylated upon invasion may play an important role in host defence. Proteomic studies interrogating the SUMOylome after infection with different pathogens may lead to the identification of these proteins. Our results revealed that the CPS and LPS O-polysaccharide mediated the *Klebsiella*-triggered decrease in SUMOylation in macrophages, in a TLR4-dependent manner. This is in perfect agreement with previous findings demonstrating that TLR4 recognises both *K. pneumoniae* polysaccharides [13,29]. The conventional wisdom indicates that TLR signalling is crucial for host defence whereas our findings demonstrate that pathogens can evolve to hijack TLR-governed signalling to attenuate host defences, thus revealing a new angle in the host-pathogen arms race. Our findings are conceptually different from previously described antagonistic strategies of TLR-signalling by pathogens to avoid activation of innate immune receptors [50]. We have recently demonstrated that *Klebsiella* targets another pathogen recognition receptor, NOD1, to attenuate the activation of NF-κB-controlled inflammation suggesting that manipulation of pathogen recognition receptors is one of the key features of *K. pneumoniae* anti-immune strategies [15]. Collectively, we put forward the notion that hijacking the activation of immune receptors to promote virulence is an emerging theme in the infection biology of pathogens. Providing further support to this notion, Arpaia and co-workers have recently shown that *S. typhimurium* requires cues from TLR-controlled signalling to regulate virulence genes necessary for intracellular survival, growth, and systemic infection [51].

A common theme for the battle of the host SUMO system versus bacterial pathogens emerges from the discoveries of this work and other recent publications investigating the interplay between SUMOylation and *Listeria, Salmonella* and *Shigella* [5–8,49]. The message is that regardless of the molecular mechanisms employed to reduce SUMOylation the outcome is identical, i.e. the global decrease of SUMO-conjugated proteins. Future studies are warranted to identify the set of SUMOylated proteins playing a crucial role in host defence against *Klebsiella*, and whether these are common to the other infections. It is tempting to consider that inhibition of deSUMOylation strategies could be a viable strategy to boost human defence mechanisms. This is challenging, as the evidence suggests that each pathogen uses different molecular mechanisms to abrogate SUMOylation, making it difficult to develop broad-spectrum therapeutics. In turn, an intriguing and exciting option to us is to develop drugs that increase SUMOylation. Targeting SUMOylation is already being considered to develop new therapeutics against cancer and heart failure [52–54]. Therefore, and based on the results shown here, it is well worth exploring the option of testing such drugs to treat bacterial infections.

## MATERIALS AND METHODS

### Cell culture

Lung epithelial A549 cells (ATCC CCL-185), murine alveolar macrophages MH-S (ATCC CRL-2019) and human monocytes THP-1 (ATCC TIB-202) were grown in Roswell Park Memorial Institute (RPMI) 1640 Medium (Gibco 21875) supplemented with 10% heat-inactivated fetal calf serum (FCS), 10 mM HEPES (Sigma), 100 U mL^-1^ penicillin, and 0.1 mg mL^-1^ streptomycin (Gibco) at 37 °C in a humidified 5% CO_2_ incubator.

Human airway epithelial cell line, NuLi-1 (ATCC CRL-4011), was grown in Bronchial Epithelial Cell Basal Medium (BEBM, Lonza CC-3171) and BEGM SingleQuot Kit Supplements and Growth Factors (Lonza CC-4175). Flasks and plates were coated with collagen type IV from placenta (Sigma C-7521).

Immortalised BMDM (iBMDM) from WT, *TLR4*^-/-^, *MyD88*^-/-^, *TRAM/TRIF*^-/-^ mice on a C57BL/6 background, were obtained from BEI Resources (NIAID, NIH) (repository numbers NR-9456, NR-9458, NR-15633 and NR-9568, respectively). iBMDM cells were grown in Dulbecco’s Modified Eagle Medium (DMEM; Gibco 41965) supplemented with 10% heat-inactivated FCS, 100 U mL^-1^ penicillin, and 0.1 mg mL^-1^ streptomycin (Gibco) at 37 °C in a humidified 5% CO_2_ incubator.

Cells were routinely tested for *Mycoplasma* contamination.

To isolate BMDMs (Bone Marrow-Derived Macrophages), tibias and femurs from C57BL/6 or *IFNAR1*^*-/-*^ mice (C57BL/6 background) were removed using a sterile technique and the bone marrow was flushed with fresh medium. To obtain macrophages, cells were plated in DMEM medium supplemented with filtered L929 cell supernatant (a source of M-CSF) and maintained at 37 °C in a humidified atmosphere of 5% CO_2_ for 4 days. Medium was replaced with fresh supplemented media after one day.

### Ethics statement

The experiments involving mice were approved by the Queen’s University Belfast’s Ethics Committee and conducted in accordance with the UK Home Office regulations (Project Licence PPL2700). Female C57BL/6 mice (8–9 weeks of age) were mock-infected with PBS (*n*=4) and infected with the wild-type strain (*n*=4). Animals were randomised for interventions, but researchers processing the samples and analysing the data were aware which intervention group corresponded to which cohort of animals.

### Intranasal murine infection model

Female mice were infected intranasally with ∼3×10^5^ Kp52145 in 30 μL PBS (*n*=4). Control mice were inoculated with 30 μL sterile PBS (*n*=4). After 48 h, mice were euthanised using a Schedule 1 method according to UK Home Office-approved protocols. Right lung samples from infected and uninfected control mice were immersed in 1 mL of sterile PBS on ice. Samples were homogenised using a VDI 12 tissue homogeniser and protein was quantified using a bicinchoninic acid (BCA) Protein Assay Kit (Thermo Scientific). 100 μg of total lung protein were lysed in 2x sample buffer (4% [w/v] SDS, 10% [v/v], 2-mercaptoethanol, 20% [w/v], glycerol, 0.004% [w/v], bromophenol blue, 0.125 M Tris-HCl pH 6.8), sonicated for 10 seconds at 10% amplitude (Branson Sonifier), boiled at 95 °C for 5 min and centrifuged at 12000 x *g* for 1 min. 50 μg of total lung protein were resolved by standard 8% SDS-PAGE and electroblotted onto nitrocellulose membranes for immunoblotting.

### Bacterial strains

Kp52145 is a clinical isolate (serotype O1:K2) previously described [47,55]. 52145-Δ*wca*_*K2*_ is an isogenic mutant of Kp52145 lacking capsule that has been previously described [56]. To generate the 52145-Δ*glf* mutant lacking the O-antigen of LPS, and the 52145-Δ*wca*_*K2*_ Δ*glf* mutant, a 904 bp product from the centre of the *glf* gene was amplified using primers Kp52_glf_F2 (5’-GCATGAATTCATGGTACATGTCTATGGACC-3’) and Kp52_glf_R2 (5’-GCATGAATTCCATCCATATCAAGGTAACGG-3’) and digested with *Eco*RI. This was then ligated into plasmid pSF100 which had been linearised with *Eco*RI. The resulting plasmid pSF100-*glf* was transformed into *E. coli* β2163 and mobilised by conjugation into Kp52145 and 52145-Δ*wca*_*K2*_, and integrated into the chromosome, disrupting the *glf* gene. The resulting integrants were selected by growth on LB plates supplemented with carbenicillin at 50 μg mL^-1^. Chromosomal integration of the plasmid was confirmed by PCR, and loss of the O-antigen was confirmed by LPS extraction and SDS-page analysis.

### Infection conditions

Bacteria were grown in 5 mL LB at 37 °C, harvested at exponential phase (2500 x *g*, 20 min) and adjusted to an optical density of 1.0 at 600 nm in PBS (5×10^8^ CFU mL^-1^). Infections were performed using a multiplicity of infection (MOI) of 100 bacteria per cell. Infections were performed in the same media used to maintain the cell-line without antibiotics and incubated at 37 °C in a humidified 5% CO_2_ incubator. THP-1 monocytes were differentiated into macrophages using phorbol 12-myristate 13-acetate (PMA) at 5 ng mL^-1^ at the time of seeding and infections were performed two days later. Infections of MH-S and iBMDMs were performed the day after seeding. In all macrophage infections extending beyond 1 h, the medium was replaced after 1 h with fresh medium containing 100 µg mL^-1^ Gentamicin (Sigma) to kill extracellular bacteria. To synchronise infections, plates were centrifuged at 200 x *g* for 5 min. Infections of epithelial cells (A549 and NuLi-1) were performed two days after seeding with starvation for 16 h before infection using RPMI 1640 Medium (Gibco 21875) supplemented only with 10 mM HEPES.

### Transfection conditions

Transfection of A549 cells with siRNAs or plasmids was carried out using Lipofectamine 2000 (Invitrogen) lipofection reagent according to manufacturer’s instructions. For transfection of siRNAs, 1.2×10^5^ cells (12-well plate) were transfected in suspension with 20 nM siRNA using 2 μL of Lipofectamine 2000 in a final volume of 1 mL. Infections were carried 48 h post transfection.

All siRNA duplexes used for *in vitro* studies were chemically synthesised by Dharmacon (GE Healthcare). The following siRNAs sense sequences were used: h*SENP1* (5′-GUG AAC CAC AAC UCC GTA UUC-3′ [57]), h*SENP2* (5′-GGG AGU GAU UGU GGA AUG UTT-3′ [57]), h*SENP3* (5′-GCU UCC GAG UGG CUU AUA ATT-3′ [57]), h*SENP5* (5′-GUC CAC UGG UCU CUC AUU ATT-3′ [57]), h*SENP6* (5′-GAC UUA ACA UGU UGA GCA ATT-3′ [57]), h*SENP7* (5′-CAA AGU ACC GAG UCG AAU AUU-3′ [57]), h*CSN5* (5′-GGA UCA CCA UUA CUU UAA GTT-3′ [58]), h*SKP1* (5’-GGA AGA UUU GGG AAU GGA U-3’), h*CUL1* (5’-UUG UGC CUA CCU CAA UAG AUU UU-3’), h*RBX1* (5’-AACUGUGCCAUCUGCAGGAACAA-3’), h*βTrCP* (5’-GUG GAA UUU GUG GAA CAU C-3’), h*SKP2* (5’-ACU CAA GUC CAG CCA UAA G-3’). An AllStars (AS) Negative Control scrambled siRNA (Qiagen) with no homology to any known mammalian gene was used as a negative control. Efficiency of transfection was confirmed by qPCR analysis of duplicate samples from three independent transfections by normalising to the glyceraldehyde 3-phosphate dehydrogenase (h*GAPDH*) gene and comparing gene expression in the knockdown sample with the AllStars negative control. Primers used are listed in Supplementary Table S1. The percentage of knockdown is presented in Supplementary Figure S7.

For transfection of plasmids in A549, 5×10^4^ (24-well) or 1.2×10^5^ (12-well) cells were seeded and the next day transfected with 2 μL (24-well) or 4 μL (12-well) of Lipofectamine 2000 and 1 (24-well) or 2 μg (12-well) of plasmid in a final volume of 0.5 or 1 mL, respectively. For transfection of plasmids in MH-S, 5.0×10^5^ (12-well) cells were seeded and the next day transfected with 4 μL of Lipofectamine 2000 and 2 μg of plasmid in a final volume of 1 mL. Experiments using plasmid-transfected cells were carried out 24 h post transfection.

The following plasmids were used: pcDNA3 (Invitrogen), pSUMO1 (pcDNA3-HA-SUMO1 was a gift from Junying Yuan, Addgene plasmid # 21154), pSENP2-GFP (pEGFP-C2 SENP2 was a gift from Mary Dasso, Addgene plasmid # 13382), pSENP2-FLAG (FLAG-SENP2 was a gift from Edward Yeh, Addgene plasmid # 18047), GSK3β-WT (HA GSK3 beta wt pcDNA3 was a gift from Jim Woodgett, Addgene plasmid # 14753), GSK3β-S9A (HA GSK3 beta S9A pcDNA3 was a gift from Jim Woodgett, Addgene plasmid # 14754). Plasmids were purified from a host *E. coli* C600 strain using an Endofree Maxi-Prep kit from Qiagen (QIAGEN, catalog number 12362) following the manufacturer’s recommendations.

### Immunoblot analysis

Macrophages were seeded in twelve-well plates (5.0×10^5^ cells per well) and grown for 24 h. A549 were seeded in twelve-well plates (1.2×10^5^ cells per well) and grown for 48 h prior to infection. Cells were infected with *K. pneumoniae* strains for different time points as indicated in the figure legends. Cells were then washed in 1 mL of ice-cold PBS and lysed in 80 μL of 2x sample buffer. The cell lysates were sonicated for 10 seconds at 10% amplitude (Branson Sonifier), boiled at 95 °C for 5 minutes and centrifuged at 12000 x *g* for 1 min. 20% of the cell lysates were resolved by standard 8 or 12% SDS-PAGE and electroblotted onto nitrocellulose membranes. Membranes were blocked with 4% bovine serum albumine (w/v) in TBST and protein bands were detected with specific antibodies using chemiluminescence reagents and a G:BOX Chemi XRQ chemiluminescence imager (Syngene) or using fluorescent antibodies and imaging in a Li-cor Odyssey model 9120.

The following antibodies were used: anti-SUMO1 (1 µg, Sigma S8070),, anti-SUMO2/3 (1:1000, Santa Cruz Biotechnology sc-32873), anti-Ubc9 (1:1000, Santa Cruz Biotechnology sc-10759), anti-SENP2 (1:1000, Santa Cruz Biotechnology sc-67075), anti-K48 Polyubiquitin (Cell Signaling 4289), anti-Cullin-1 (1:200, Santa Cruz Biotechnology sc-12761), anti-Flag M2 (1 μg, Sigma F3165), anti-HA (1:1000, Santa Cruz Biotechnology sc-805), anti-Sae1 (1:1000, Sigma Aldrich WH0010055M1), anti-Sae2 (1:1000, Sigma Aldrich SAB1306998), anti-RanGAP1 (1:1000, Santa Cruz Biotechnology sc-28322).

Immunoreactive bands were visualized by incubation with horseradish peroxidase-conjugated goat anti-rabbit immunoglobulins (1:5000, BioRad 170-6515) or goat anti-mouse immunoglobulins (1:5000, BioRad 170-6516).

To detect multiple proteins, membranes were reprobed after stripping of previously used antibodies using a pH 2.2 glycine-HCl/SDS buffer. To ensure that equal amounts of proteins were loaded, blots were reprobed with mouse anti-human tubulin (1:3000, Sigma T5168). SUMO1 smears above 90 kDa (excluding therefore the high intensity RanGAP1 bands present at 80 kDa) were quantified using Image Studio Lite version 5.2 (Li-cor) and the modification normalised to α-Tubulin. Graphs represent fold change compared to untreated non-infected cells as indicated in figure legends.

### Co-immunoprecipitation analysis

Cells were seeded (3×10^5^ cells per well) in six-well plates and two wells were used per sample. At the designated time points, cells were washed with pre-chilled PBS (1 mL) and then lysed with 500 μL of pre-chilled RIPA buffer (1% [w/v] Triton X-100, 1% [w/v] Sodium Deoxycholate, 0.1% [w/v] SDS, 0.15 M NaCl, 50 mM Tris-HCl, pH 7.2, 1 mM PMSF and Halt Protease Inhibitor Cocktail (Thermo Scientific)) for 30 min on ice. Lysates were centrifuged at 12000 x *g* for 15 min at 4 °C. Supernatants were transferred to pre-chilled tubes and incubated for 2 h with the appropriate antibody (1 μg) at 4 °C with rocking. This was followed by the addition of protein A/G agarose beads (20 μL per sample; Santa Cruz Biotechnology, sc-2003) and incubation overnight at 4 °C with rocking. Immunoprecipitates were collected by centrifugation at 1000 x *g* for 2 min at 4 °C and the beads were then washed two times with RIPA buffer (500 μL) lacking the protease inhibitor mixture. The beads were resuspended in 2x sample buffer (40 μL) and boiled for 5 min at 95 °C. Samples were centrifuged at 12000 x *g* for 1 min and subjected to immunoblotting. The immunoreactive bands were detected by incubation with Clean-Blot IP Detection Reagent (1:1000, Thermo Scientific 21230).

### Nuclear fractionation

A549 cells were seeded in six well tissue culture plates at a density of 3×10^5^ cells per well and infected after 2 days at a MOI of 100:1. Cells were collected in 500 µL of PBS and spun for 4 min at 1000 x *g* at 4 °C. The pellet was resuspended in 100 µL of cytoplasmic extract buffer (10 mM HEPES, 60 mM KCl, 1 mM EDTA, 0.075% [w/v] Triton X-100, 1 mM PMSF, 1 mM DTT and Halt protease inhibitor cocktail, pH 7.6) and incubated on ice for 4 minutes. The mixture was then spun at 1000 x *g* for 4 min at 4 °C. The cytoplasmic extract corresponding to the supernatant fraction was mixed with sample buffer (5x). The pellet was washed in 100 µL of cytoplasmic buffer without Triton X-100 and resuspended in 100 µL of sample buffer (2x).

### Intracellular survival determination

MH-S cells were seeded in twelve-well plates at a density of 5×10^5^ cells per well and infected the following day with a MOI of 100:1. Plates were incubated at 37 °C in a humidified 5% CO_2_ atmosphere. After 30 min contact, cells were washed with PBS and incubated for additional 150 min with 1 mL RPMI 1640 containing 10% FCS, 10 mM HEPES and gentamicin (100 μg mL^-1^) to eliminate extracellular bacteria. To determine intracellular bacterial load, cells were washed twice with PBS and lysed with 300 μL of 0.1% (w/v) Saponin in PBS for 5 min at 37 °C. Serial dilutions were plated on LB to quantify the number of intracellular bacteria. Bacterial adhesion was determined by serial dilutions after lysis with saponin as described before at the time of contact (30 min). Bacterial load is represented as CFU per well. All experiments were carried out with duplicate samples on at least five independent occasions.

### Microscopy

A549 cells were seeded (5×10^4^ cells per well) in 24-well plates with sterile round glass cover slips and grown for 24 h to 70-80% confluence. Cells were transfected using Lipofectamine 2000 with plasmids encoding pEGFP-C2 SENP2 (gift from Mary Dasso, Addgene plasmid # 13382) and infected 24 h later. After 5 h incubation with either *K. pneumoniae* or PBS (control), medium was removed, and the cells were gently washed twice with PBS. Cells were then fixed by the addition of 2% (w/v) paraformaldehyde (250 μL) for 20 min. Cells were washed twice with PBS and kept at 4 °C in PBS supplemented with 1 mM NH_4_Cl until staining. Nuclei were stained with Hoechst 33342 (1.5 μg mL^-1^; Sigma) for 2 h, washed with PBS and were mounted with ProLong Gold Antifade Mountant (Molecular Probes Inc.). Fluorescence images were captured using the ×40 objective lens on a Leica DM5500 microscope equipped with the appropriate filter sets. Acquired images were analysed using the LAS imaging software (Leica).

### MicroRNA analysis

Cells were washed in PBS and miRNA was extracted using the mirPremier microRNA Isolation Kit (Sigma) according to the manufacturer’s instructions. Complementary DNA was generated from 100 ng of miRNA enriched RNA using the NCode VILO miRNA cDNA Synthesis kit (Invitrogen) following manufacturer’s recommendations. Samples were assayed by quantitative real-time PCR using KAPA SYBR FAST (KAPA Biosystems) and Stratagene Mx3005P qPCR System (Agilent Technologies). Primers were designed using the software miRprimer [59] and are listed in Supplementary Table 1. Thermal cycling conditions were as follows: 50 °C for 2 min for UDG incubation, 95 °C for 3 min for enzyme activation, 42 cycles of denaturation at 95 °C for 15 s and 1 min annealing and amplification at 60 °C, followed by melt curve analysis (1 min at 95 °C, 30 s at 60 °C ramping to 30 s at 95 °C). Each cDNA sample was tested in duplicate and relative mRNA quantity was determined by the comparative threshold cycle (ΔΔC_T_) method using *snoRNA-202* miRNA gene normalisation.

For inhibition of the specific miRNAs in cells, 20 nM of the corresponding antagomir were transfected using Lipofectamine RNAiMax (Invitrogen) 24 hours before infection at the time of macrophage seeding (5×10^5^ cells per well). Mouse miRIDIAN microRNA hairpin inhibitors were purchased from Dharmacon: negative control (catalog number IN-001005-01-05), mmu-let-7b-5p (catalog number IH-310504-07-0002), mmu-let-7f-5p (catalog number IH-310509-08-0002), mmu-let-7g-5p (catalog number IH-3100374-08-0002), mmu-let-7i-5p (catalog number IH-310375-08-0002). Efficiency of transfection was confirmed by qPCR analysis of duplicate samples by comparing gene expression with the negative control. Samples were collected as described above 24 h post-transfection, from three independent transfections and is presented in Supplementary Figure S7.

### RNA extraction and quantitative real-time PCR analysis

Cells were washed twice in PBS and RNA was extracted using TRIzol Reagent (Ambion) according to the manufacturer’s instructions. Duplicate cDNA preparations from each sample were generated from 1 μg of RNA using Moloney murine leukaemia virus (M-MLV) reverse transcriptase (Sigma-Aldrich) according to the manufacturer’s instructions. Quantitative real-time PCR analysis of gene expression was undertaken using the KAPA SYBR FAST qPCR Kit and Stratagene Mx3005P qPCR System (Agilent Technologies). Thermal cycling conditions were as follows: 95 °C for 3 min for enzyme activation, 40 cycles of denaturation at 95 °C for 10 s and annealing at 60 °C for 20 s. Primers used in qPCR reactions are listed in Supplementary Table 1. cDNA samples were tested in duplicate and relative mRNA quantity was determined by the comparative threshold cycle (ΔΔC_T_) method, using hypoxanthine phosphoribosyltransferase 1 (m*HPRT*) gene normalisation for mouse samples or the glyceraldehyde 3-phosphate dehydrogenase (h*GAPDH*) gene for human samples.

### Quantification of cytokines

Infections were performed in twelve-well plates (1.2×10^5^ cells per well for A549 and 5.0×10^5^ cells per well for MH-S) using Kp52145 at a MOI of 100:1. Supernatants from infected cells were collected at the time points indicated in the figure legends and spun down at 12000 x *g* for 5 min to remove any debris. TNFα in the supernatants was determined using a Murine TNFα Standard TMB ELISA Development Kit (PeproTech, catalog number 900-T54) and a Human IL-8 Standard TMB ELISA Development Kit (PeproTech, catalog number 900-T18), according to manufacturer’s instructions. Experiments were run in duplicate and repeated at least three times.

For quantification of type-I IFN (INFα/β) in the supernatants of iBMDMs, cells were seeded (12-well plate, 5×10^4^ cells per well) and grown for 24 h. Cells were then infected (MOI 100:1) for 3 h, and supernatants were collected. Murine type-I IFN were detected using the B16-Blue IFN-α/β reporter cells (Invivogen) which carry a SEAP reporter gene under the control of the IFN-α/β-inducible ISG54 promoter and that have an inactivation of the IFN-γ receptor. Briefly, supernatants from iBMDM infections were incubated with the reporter cell line and levels of SEAP in the supernatants were determined using the detection medium QUANTI-Blue (Invivogen) after 24 h as per manufacturer’s instructions using recombinant mouse IFNβ (PBL assay science, catalog number 12401-1) as a standard.

### Statistical analysis

Statistical analyses were performed using ANOVA, or when the requirements were not met, by unpaired t-test. Multiple comparison was performed using Holm-Sidak’s multiple comparisons test. P-values of < 0.05 were considered statistically significant. Normality and equal variance assumptions were tested with the Kolmogorov–Smirnov test and the Brown– Forsythe test, respectively. All analyses were performed using GraphPad Prism for Windows (version 5.03) software.

## ACKNOWLEDGEMENTS

We thank the members of the JAB laboratory for their thoughtful discussions and support with this project. We thank Daniel Longley for kindly providing us with the SCF siRNAs Skp1, Cul1, Rbx1, Skp2 and βTrCP used in this study. LH is the recipient of a Queen’s University Belfast Research Fellowship. KP and FNV are fellows funded by European Union Seventh Framework Programme Marie Curie Initial Training Networks (FP7-PEOPLE-2012-ITN) for the project INBIONET under grant agreement PITN-GA-2012-316682. This work was supported by Marie Curie Career Integration Grant U-KARE (PCIG13-GA-2013-618162); Biotechnology and Biological Sciences Research Council (BBSRC, BB/L007223/1, and BB/P006078/1) and Queen’s University Belfast start-up funds to JAB.

## AUTHORS CONTRIBUTIONS

JS-P and JAB conceived the study and wrote the first draft of the manuscript. JS-P, KP, FNV, AD, CGF and LH performed the experiments and contributed data for this work. JS-P, KP, FNV, AD, CGF, LH and JAB contributed to and approved the final version of the manuscript.

## SUPPLEMENTARY TABLE LEGENG

**Table S1. List of primers used in this work**.

## SUPPLEMENTARY FIGURE LEGENDS

**Figure S1 – *Klebsiella pneumoniae* decreases SUMO-conjugated proteins in NuLi-1 epithelial cells**. Immunoblot analysis of SUMO1 and tubulin levels in lysates of NuLi-1 cells infected with Kp52145 for the indicated times. Data is representative of at least three independent experiments. n.i. – non-infected control. SUMO1 smears were quantified from three independent experiments using Image Studio Lite (Li-cor) and the modification normalised to α-Tubulin. The graph represents fold change compared to control non-infected cells. *P ≤ 0.05; versus n.i. determined using one way-ANOVA with Bonferroni’s multiple comparisons test.

**Figure S2 – Role of deSUMOylases in *K. pneumoniae*-induced decrease in SUMO-conjugated proteins**.

A. Immunoblot analysis of SUMO1 and tubulin levels in lysates of control (AS), Senp3, Senp5, Senp6 or Senp7 siRNA-transfected A549 cells infected with Kp52145 for 5 h. AS-AllStars control, non-silencing siRNA. Data are representative of at least three independent experiments. SUMO1 smears were quantified from three independent experiments using Image Studio Lite (Li-cor) and the modification normalised to α-Tubulin. The graph represents fold change compared to control A.S.-transfected non-infected cells. ***P ≤ 0.001;

*P ≤ 0.05 versus n.i. determined using one way-ANOVA with Bonferroni’s multiple comparisons test.

B.*senp* mRNA levels, assessed by qPCR, in A549 cells left untreated (n.i.) or infected for 5 h with Kp52145. Values are presented as the mean ± SD of three independent experiments measured in duplicate.

**Figure S3 – *K. pneumoniae*-induced decrease in macrophage SUMO1-conjugated proteins is dependent on live bacteria**. Immunoblot analysis of SUMO1 and tubulin levels in lysates of MH-S cells infected with Kp52145 live, heat-killed for 30 min at 60 °C or UV-killed for 30 min at 10000 J, for 3 h. Data is representative of at least three independent experiments. n.i. – non-infected control. SUMO1 smears were quantified from three independent experiments using Image Studio Lite (Li-cor) and the modification normalised to α-Tubulin. The graph represents fold change compared to control non-infected cells. *P ≤ 0.05 versus n.i. determined using one way-ANOVA with Bonferroni’s multiple comparisons test.

**Figure S4 – There is no change in Ubc9, Senp2 or Cullin-1 NEDDylation in MH-S cells infected with *K. pneumoniae***.

A. Immunoblot analysis of Ubc9 and tubulin levels in lysates of MH-S cells infected with Kp52145 for the indicated times.

B. Immunoblot analysis of Senp2, Lamin A/C and tubulin levels in nuclear or cytosolic extracts of MH-S cells infected with Kp52145 for 3 h or left uninfected (n.i.).

C. Immunoblot analysis of Cullin-1 and tubulin levels in lysates of MH-S cells infected with Kp52145 for 3 h or left uninfected (n.i.).

In all panels data are representative of at least three independent experiments.

**Figure S5 – Interferon signalling is upregulated in MH-S cells infected with *K. pneumoniae* and interferon induces a decrease in SUMO1-conjugated proteins**.

A. mRNA levels of the interferon stimulated genes *isg15* (Interferon-Stimulated Gene 15), *ifit1* (Interferon Induced Protein With Tetratricopeptide Repeats 1) and *irf7* (Interferon Regulatory Factor 7), assessed by qPCR, in MH-S cells left untreated (n.i.) or infected for 3 h with Kp52145. Values are presented as the mean ± SD of three independent experiments measured in duplicate. ***P ≤ 0.001; *P ≤ 0.02; versus n.i. determined using one way-ANOVA with Tukey’s multiple comparisons test.

B. Immunoblot analysis of SUMO1 and tubulin levels in lysates of MH-S cells infected with 52145-Δ*wca*_*K2*_ Δ*glf* for 1 h or left uninfected (n.i.), pre-treated with 5000 U of IFNβ for 30 min. Cells with no interferon pre-treatment are presented as controls. SUMO1 smears were quantified from three independent experiments using Image Studio Lite (Li-cor) and the modification normalised to α-Tubulin. The graph represents fold change compared to control non-infected cells. *P ≤ 0.05 versus n.i. determined using one way-ANOVA with Bonferroni’s multiple comparisons test.

In all panels data are representative of at least three independent experiments.

**Figure S6 – SUMO1 upregulation in A549 and MH-S cells**.

A. Immunoblot analysis of Sumo1 and tubulin levels in lysates of A549 cells transfected with either empty control plasmid (pcDNA3) or SUMO1-HA tagged plasmid (pSUMO1). Cells were infected with Kp52145 for 5 h (MOI 100:1). Data are representative of at least three independent experiments.

B. Immunoblot analysis of Sumo1 and tubulin levels in lysates of MH-S cells transfected with either empty control plasmid (pcDNA3) or SUMO1-HA tagged plasmid (pSUMO1). Cells were infected with Kp52145 for 3 h (MOI 100:1). Data are representative of at least three independent experiments.

C. Adhesion in MH-S cells transfected with either empty control plasmid (pcDNA3) or SUMO1-HA tagged plasmid (pSUMO1). Cells were infected with Kp52145 for 30 min (MOI 100:1), wells were washed and bacteria were quantified by lysis, serial dilution and viable counting on LB agar plates. Values are presented as the mean ± SD of three independent experiments measured in duplicate.

D. Adhesion in antagomir transfected MH-S cells. Cells were infected with Kp52145 for 30 min (MOI 100:1), wells were washed and bacteria were quantified by lysis, serial dilution and viable counting on LB agar plates. Values are presented as the mean ± SD of three independent experiments measured in duplicate.

**Figure S7 – Efficiency of transfection of siRNA into A549 or MH-S cells**. Efficiency of transfection presented as percent (%) of knockdown after transfection. mRNA levels of the indicated transcripts were accessed 48 h post-transfection as fold change against control non-silencing agents (AS-AllStars control, non-silencing siRNA, in the case of *senp, csn5, βTrCP, cullin1, skp1, skp2* and *rbx1* and negative control for *let-7* antagomirs) after gene normalization. Values are presented as the mean ± SD of three independent experiments measured in duplicate.

In all panels data are representative of at least three independent experiments.

